# Extracting orthogonal subject- and behavior-specific signatures from fMRI data using whole-brain effective connectivity

**DOI:** 10.1101/201624

**Authors:** Vicente Pallarés, Andrea Insabato, Ana Sanjuán, Simone Kühn, Dante Mantini, Gustavo Deco, Matthieu Gilson

## Abstract

The study of brain communication based on fMRI data is often limited because such measurements are a mixture of session-to-session variability with subject- and condition-related information. Disentangling these contributions is crucial for real-life applications, in particular when only a few recording sessions are available. The present study aims to define a reliable standard for the extraction of multiple signatures from fMRI data, while verifying that they do not mix information about the different modalities. In particular, condition-specific signature should not be contaminated by subject-related information. Practically, signatures correspond to subnetworks of directed interactions between brain regions (typically 100 covering the whole brain) supporting the subject and condition identification for single fMRI sessions. The key for robust prediction is using effective connectivity instead of functional connectivity. Our method demonstrates excellent generalization capabilities for subject identification in two datasets, using only a few sessions per subject as reference. Using another dataset with resting state and movie viewing, we show that the two signatures related to subjects and tasks correspond to distinct subnetworks, which are thus topologically orthogonal. Our results set solid foundations for applications tailored to individual subjects, such as clinical diagnostic.

## 1 Introduction

Blood-oxygen-level dependent (BOLD) signals in functional magnetic resonance imaging (fMRI) have been used for more than two decades to observe human brain activity and relate it to cognitive functions [Cordes et al., 2000, Amunts et al., 2014]. Even at rest, the brain exhibits patterns of correlated activity between distant areas [Biswal et al., 1995, Raichle et al., 2001]. Although a consensus has yet to be reached, many methods have been proposed to quantify this activity and extract functionally-relevant information, beyond the widely-used functional connectivity (FC) that measures the statistical dependencies between the BOLD activities of brain regions. For instance, interest in the temporal BOLD structure for both individual regions [He, 2011] and between areas (via the cross-covariance lags at the scale of seconds) [Mitra et al., 2015] has grown; the ‘dynamic FC’ was defined to quantify the BOLD correlations at the scale of minutes [Preti et al., 2017, Gonzalez-Castillo and Bandettini, 2017]. Following fundamental discoveries about brain functions, fMRI has increasingly been used to complement clinical diagnostic for neuropathologies [Matthews and Hampshire, 2016]. Resting-state fMRI has also been found to be informative about neuropsychiatric disorders [Greicius, 2008]: alterations in FC correlate with and can predict the clinical scores of several diseases [Kurth et al., 2015, Rahim et al., 2017]. This defines “signatures” in the form of subnetworks of links between brain regions that consistently covary with respect to behavioral or pathological conditions.

It has recently been shown that fMRI signals are strongly biased by individual traits [Miranda-Dominguez et al., 2014, Finn et al., 2015, 2017, Calhoun et al., 2017], which should be taken into account when extracting task-specific signatures [Xie et al., 2017]. Generalizing, the mixture of session-to-session, subject-specific and condition-related variability in FC is a severe limitation for real-life applications when only a few sessions per subject can be recorded [Varoquaux et al., 2017, Woo et al., 2017]. In that respect, FC measures obtained from successive recording sessions have been found not to be reliable individually (i.e., at the level of single links), but only collectively [Shehzad et al., 2009, Mueller et al., 2015, Chen et al., 2015, Pannunzi et al., 2017]. Because previous studies [Finn et al., 2015, Miranda-Dominguez et al., 2014, Finn et al., 2017, Xie et al., 2017] were limited to datasets with at most 3 sessions for the same behavioral condition per subject, we aim to rigorously assess the generalization capability of prediction methods to future (unseen) data. Meanwhile, we examine how multivariate classification can disentangle signatures for distinct modalities.

Distributed signatures in FC across the whole brain have been observed in memory tasks [Rissman and Wagner, 2012] or when the subject experiences psychological pain [Chang et al., 2015]. Moreover, the etiology of many mental disorders is unknown: They are suspected to arise from network dysfunction, as reported for large-scale FC alterations in patients with schizophrenia [Hoptman et al., 2012]. These examples strongly point in favor of whole-brain approaches to study high-level cognition [Deco et al., 2011] and brain diseases [Deco and Kringelbach, 2014]; in contrast, focusing on a few cortical areas only to test hypotheses [Goebel et al., 2003, Bastos-Leite et al., 2015] may not capture sufficient information or network effects. Such whole-brain approaches typically involve a large number of parameters to estimate, which may impair the robustness of classification based on these parameters. One aim of the present study is to provide a practical answer to this trade-off.

The idea underlying the study of FC—in the broad sense—is that it reflects how brain areas dynamically bind to exchange and process information [Fries, 2005, Hipp et al., 2011, Betti et al., 2013]. To go beyond a phenomenological description of FC, we employ a large-scale effective connectivity (EC) relying on a model inversion [Gilson et al., 2016] to decompose FC into changes in network connectivity and local fluctuating activity. We borrow the EC terminology to describe interactions between brain regions from dynamic causal modeling [Friston, 2011] despite differences with our model, which will be discussed later. So far, DCM-based EC has been used for classification, but using 6 ROIs and two sessions only [Brodersen et al., 2011]. Here we stress the need for adequate quantitative methods to assess to which extent EC can predict the subject’s identity [Miranda-Dominguez et al., 2014, Frassle et al., 2015] or behavioral condition [Gonzalez-Castillo et al., 2015], because the inference of signatures—e.g., changes in FC that “simply” correlate with behavioral conditions [Li et al., 2012, Bastos-Leite et al., 2015, Gilson et al., 2017]—does not guarantee robustness in the classification performance. In other words, parameter inference hints at putative signatures, but their predictive power must be carefully verified. As mentioned earlier, the overlap between subject- and condition-related signatures may impair the classification performance, when some links are sensitive to both modalities and should be excluded (or “conditioned out” as with probabilities) to disentangle the signatures.

The present study aims to set a new standard for extracting multivariate signatures from fMRI data, here applied to discriminate between subjects or behavioral conditions. It is organized in two parts. First, we couple whole-brain EC estimation with adequate machine learning tools to control for session-to-session variability. The focus is on the comparison between EC and FC in their generalization capabilities to unseen data for the classification of single resting-state fMRI sessions with respect to (healthy) subjects [Finn et al., 2015, Miranda-Dominguez et al., 2014]. We use datasets with large numbers of sessions per subject to provide a benchmark with an unprecedented level of control. Second, we predict both subject identity and condition (rest versus movie viewing) to verify that EC can disentangle the two types of signatures. To do so, we examine the topological distribution of the EC links supporting the twofold classification, which allows us to quantify the overlap between the two signatures. This second part thus aims to assess the generalization capability of EC for multivariate classification.

## 2 Results

### 2.1 Functional and effective connectivity as measures of brain network dynamics

In this study we used fMRI data from the three datasets described in Table 1. Classical functional connectivity (corrFC) was calculated using the pairwise Pearson correlation coefficient (PCC) between the time courses of the *N* regions of interest (ROIs), obtaining an *N* × *N* symmetric matrix for each recorded session (*N* = 116 for Datasets A and B, *N* = 66 for Dataset C); see Eq. (2) in Methods. In parallel, we used the whole-brain dynamic model [Gilson et al., 2016, 2017] in Fig. 1B: Each ROI is a node in a noise-diffusion network whose topology (skeleton) is determined by the structural connectivity (SC) obtained from diffusion tensor imaging (DTI) or similar techniques. In the model, the global pattern of FC arises from the local variability Σ_*i*_ that propagates via the network connections EC_*ij*_ (from *j* to *i*). To fit each fMRI session, all relevant EC_*ij*_ and Σ_*i*_ parameters are iteratively tuned such that the model spatio-temporal FC—as measured by FC0 (0-lag covariances) and FC1 (1-lag shifted covariances), see Eq. (1) in Methods—best reproduces the empirical counterpart. A detailed description of the model and the estimation procedure is provided in Methods. In essence, the model inversion decomposes the empirical matrices (FC0,FC1) into two estimates EC and Σ, which can be seen as multivariate biomarkers for the brain dynamics in each fMRI session.

**Table 1.** Details of the datasets used in the present study.

| Dataset name | Acquisition / Reference | Subject number | Sessions per subject | Session duration |
| --- | --- | --- | --- | --- |
| Dataset A1 | Day2day project [Filevich et al., 2017] | 6 | 40-50 | 5 minutes |
| Dataset A2 | Day2day project [Filevich et al., 2017] | 50 | 1 | 5 minutes |
| Dataset B | CoRR [Zuo et al., 2014] | 30 | 10 | 10 minutes |
| Dataset C | [Mantini et al., 2012, Gilson et al., 2017] | 19 | 3 (rest) and 2 (movie) | 10 minutes |

session.

**Figure 1:**
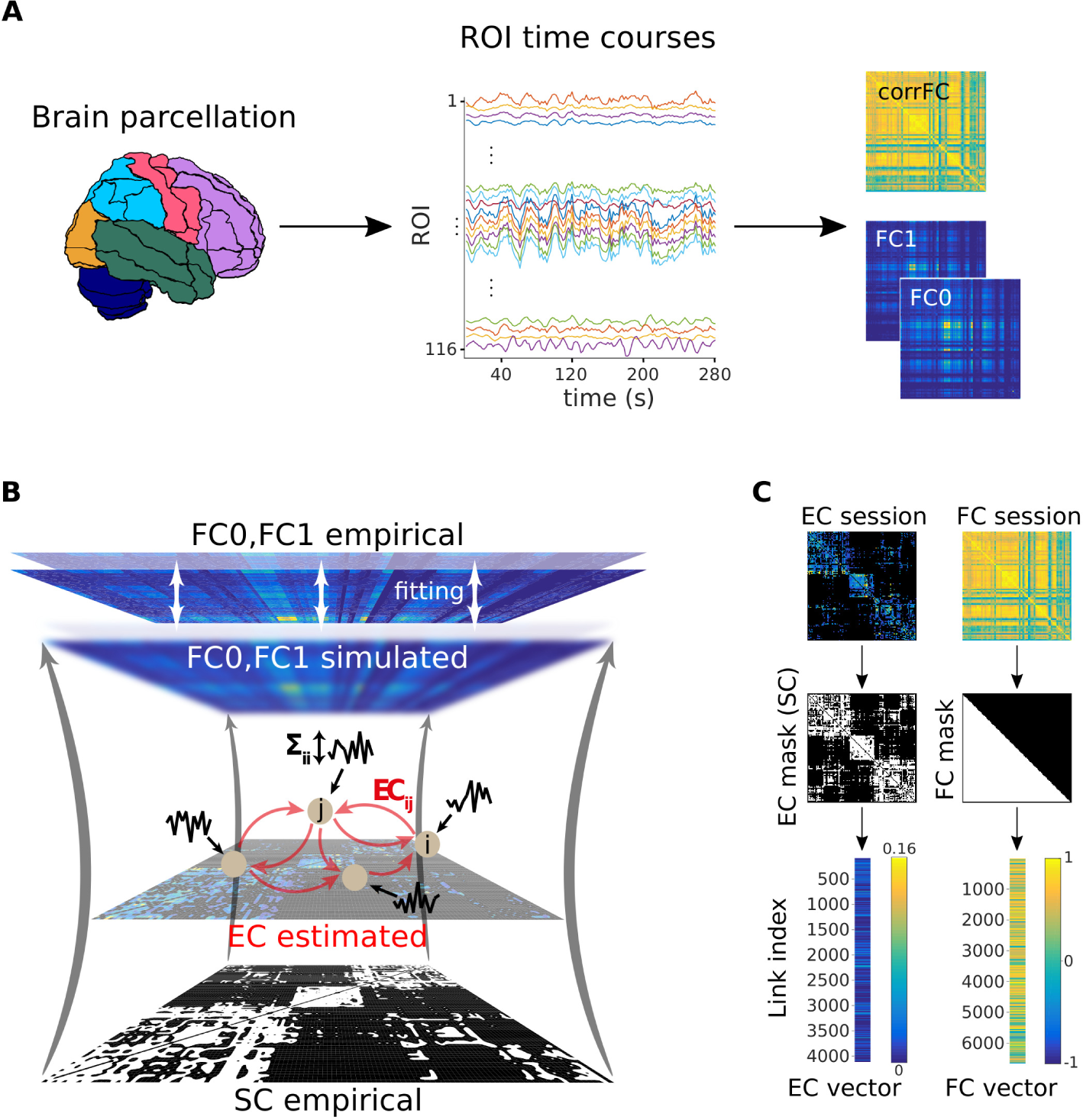
**Workflow for the calculation of the connectivity measures from fMRI measurements. A)** After a standard pre-processing pipeline, a parcellation covering the whole-brain is applied to extract BOLD time series: 116 ROIs for the AAL parcellation used here and 66 ROIs for the Hagmann parcellation, with each color representing an anatomical subsystem of several ROIs. Here we consider two versions of functional connectivity: the classical corrFC corresponding to the Pearson correlation coefficient (PCC) between pairs of time series; the spatiotemporal FC embodied by the two covariance matrices FC0 and FC1 without and with time shift, respectively. Details are provided in Methods, see Eq. (1) and Eq. (2). **B)** Whole-brain network model to extract effective connectivity (EC) from fMRI measurements. The local fluctuating activity (where Σ_*i*_ is the variance of the input to each region i) propagates via the recurrent EC to generate the correlation patterns at the network level. Structural connectivity (SC, bottom) obtained using DTI determines the skeleton of EC. The fitting procedure iteratively tunes EC and Σ such that the model best reproduces the empirical FC0 and FC1. **C)** Each corrFC matrix is symmetric and has all diagonal elements equal to 1, so that only 6670 independent links are retained for identification/classification (lower triangle). Likewise, the EC matrix has 4056 non-zero elements that are used in the classification (density of 30%).

### 2.2 Structure of individual session-to-session variability for EC and FC

Before considering classification, we examined the distributions of connectivity measures extracted from repeated scans for individual subjects. Using Datasets A and B, we compared the capability of the 4 connectivity measures (corrFC, FC0, FC1 and EC), as well as Σ, in terms of within- and between-subject similarity (WSS and BSS, respectively), as a first step toward subject identification. For each pair of sessions, the similarity *S_X_* was calculated using the PCC between two vectorized connectivity measures *X* in Fig. 1C (non-zero elements for EC, low-triangle elements for corrFC). In the matrix of *S*_EC_ values for Dataset A1 (Fig. 2A), 6 diagonal blocks with larger values corresponding to the WSS can be noticed; the remaining matrix elements correspond to BSS. Fig. 2B compares the distributions of *S*_EC_ and *S*_corrEC_: WSS and BSS distributions are better separated for EC than for corrFC. In other words, sessions from the same subject are more similar to one another, and more different from those of other subjects, viewed from the EC than the corrFC perspective. This suggests a better capability for EC to discriminate between subjects. Note that the BSS from Datasets A1 (6 subjects) and A2 (50 subjects) remarkably overlap for both corrFC and EC, showing that BSS for 6 subjects generalizes well to larger numbers.

**Figure 2:**
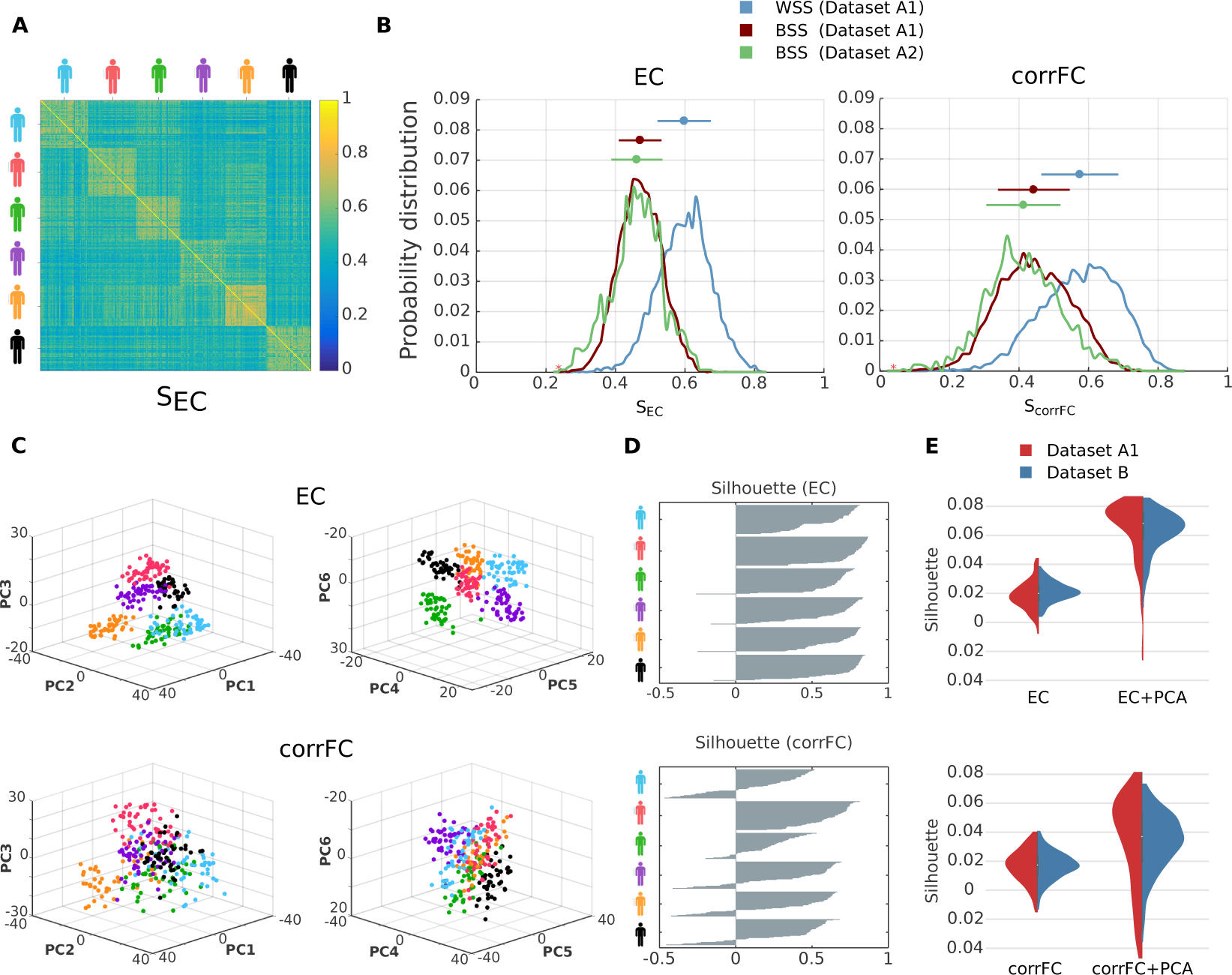
**Within- and between-subject similarity (WSS and BSS, respectively) for EC and corrFC. A)** Matrix of similarity values for EC between all pairs of sessions from Dataset A1. ECs from two sessions are transformed into two vectors (as illustrated in Fig. 1C), from which the PCC is calculated to obtain SEC (see Eq. (12) in Methods). The sessions are grouped by subjects, as indicated by the colored symbols. **B)** The left panel shows that distributions of WSS (blue) and BSS (red) values for Datasets A1—corresponding to diagonal and off-diagonal blocks in panel A, respectively—and of BSS (green) for Dataset A2. The right panel shows the corresponding distributions for corrFC. The above error bars indicate the means and standard deviations, indicating a smaller overlap between WSS and BSS for EC. **C)** Visualization of the sessions of Dataset A1 in the space of the first 6 principal components, or PCs (split into the left and right panels), obtained from PCA for EC (top row) and corrFC (bottom row). Each point corresponds to a session and each color to one of the 6 subjects, as in panel A. **D)** Silhouette coefficients of each session in panel C (see main text and Eq. (15) in Methods for further details). **E)** Distribution of the silhouette coefficients for EC (top panel) and corrFC (bottom panel): comparison between the original link space (left) and the PCA space (right, corresponding to panel D). Both Datasets A1 (6 subjects with 6 PCs, in red) and B (30 subjects with 30 PCs, in blue) are represented by the violin plots; see also Figure S3 about the choice for the number of PCs. Note the larger silhouette coefficients for EC than for corrFC.

These qualitative observations are confirmed by Table 2 that summarizes the Kolmogorov-Smirnov (KS) distance between the similarity distributions (blue versus red and blue versus green in Fig. 2B). EC gave larger KS distance than corrFC and all other FC-related measures. Note that we also calculated KS distance using only the links in corrFC and FC0 corresponding to the 4056 existing connections in EC (determined by SC), in order to compensate for the relative sparsity of EC links as compared to corrFC and FC0; however, this did not change the results. Lastly, the diagonal elements of Σ showed the smallest distances. In the following, we focus on EC and corrFC. Supplementary Figures S1 and S2 show the similarity distributions for FC0, FC1, corrFC/SC and E using Datasets A1 and B.

**Table 2.** Kolmogorov-Smirnov (KS) distance between WSS and BSS distributions.

| Measure | Description | Dataset A1 | Dataset A1 & A2 |
| --- | --- | --- | --- |
| EC | EC weights estimated with the model | 0.6440 | 0.6581 |
| corrFC | FC computed as Pearson correlation | 0.4517 | 0.5477 |
| FC0 | FC computed as 0-lag covariance matrix | 0.3685 | 0.4888 |
| FC1 | FC computed as 1-lag covariance matrix | 0.4139 | 0.5118 |
| corrFC/SC | FC as Pearson correlation and masked with SC | 0.4769 | 0.5729 |
| FC0/SC | FC as 0-lag covariance matrix and masked with SC | 0.3770 | 0.4644 |
| $\Sigma$ | Local variability for each ROI estimated with the model | 0.3778 | 0.3287 |

The high dimensionality of our connectivity measures may reduce their predictive power, known as the Hughes phenomenon [Hughes, 1968]. This is especially important in our case where the number *p* of dimensions for the multivariate measures (Dataset A1: *p* = 4056 for EC, *p* = 6670 for corrFC) exceeds the number *n* of samples (*n* = 6 subjects × 50 sessions = 300 samples). To further characterize individual variability over sessions, we performed a reduction of dimensionality using a principal component analysis (PCA) on the sessions of Dataset A1. To the naked eye, the colored clouds representing all sessions for each subject exhibit smaller overlap for EC than corrFC in Fig. 2C, where data are projected onto the first 6 principal components (PCs).

We quantified the clustering degree of these clouds using a silhouette coefficient for each session, ranging from -1 (poor clustering) to 1 (perfect clustering); see Eq. (15) in Methods. As shown in Fig. 2D, EC produced larger (almost all positive) silhouette values than corrFC, confirming the visual impression of Fig. 2C. Silhouette coefficients were calculated on the data projected onto the first 6 PCs, i.e. the number of PCs that maximized the silhouette coefficient (see Supplementary Figure S3). As can be seen in Fig. 2E, the silhouette coefficients for the data in the original link space (left violin plots) are smaller than those for data in the PCs space (right); for Dataset B (in blue), the first 30 PCs are used. Consequently, PCA may facilitate the identification of subjects by reducing the dimensionality of the data, but this has to be properly tested.

### 2.3 Subject identification using EC is more robust than using FC

Now we turn to the main goal of our study: the classification of single sessions based on EC or corrFC. In this section, we start with attributing sessions to subjects. Subject identification for a large cohort (∼ 100 subjects) was recently pioneered [Finn et al., 2015], relying on a k-nearest-neighbor (*k*NN) classifier with k=1 and PCC as metric. In order to classify a target session, the PCC between the target and 1 known reference session for each subject (called database) is calculated; the predicted identity for the target is that of the subject corresponding to the closest (most similar) session (see Supplementary Figure S4 and Methods for details). In contrast with previous studies using 1NN [Finn et al., 2015, 2017, Kaufmann et al., 2017], our method relies on a multinomial logistic regression (MLR) classifier, a classical tool in machine learning. MLR uses a linear model to predict the probability that an input sample belongs to a class (subject here). A technical comparison of both approaches is further detailed in Methods.

In classification algorithms the problem of overfitting describes the situation where the algorithm performs very well with the data it is trained with, but fails to generalize to new samples. Due to the high dimensionality of the connectivity measures [Hughes, 1968], it is essential to control for overfitting with an appropriate training and test procedure. Our train-test procedure and the use of large test-retest datasets—unlike previous studies [Finn et al., 2015, 2017, Vanderwal et al., 2017]—aims to provide a trustworthy characterization of the quality of the classifiers. Fig. 3A describes the train-test procedure for the identification of subjects: 1) fMRI sessions (EC in the figure) are randomly split in training and test datasets; 2) after preprocessing (orange arrows) involving within-session z-score (see Eq. (16) in Methods) followed—or not—by PCA, the classifier is optimized as illustrated for the MLR with boundaries that best predict the training dataset; 3) test set is used to verify the generalization capability of the classifier (blue arrows), by measuring to which extent the classifier boundaries, estimated with the train set, correctly classify single sessions from the test set.

**Figure 3:**
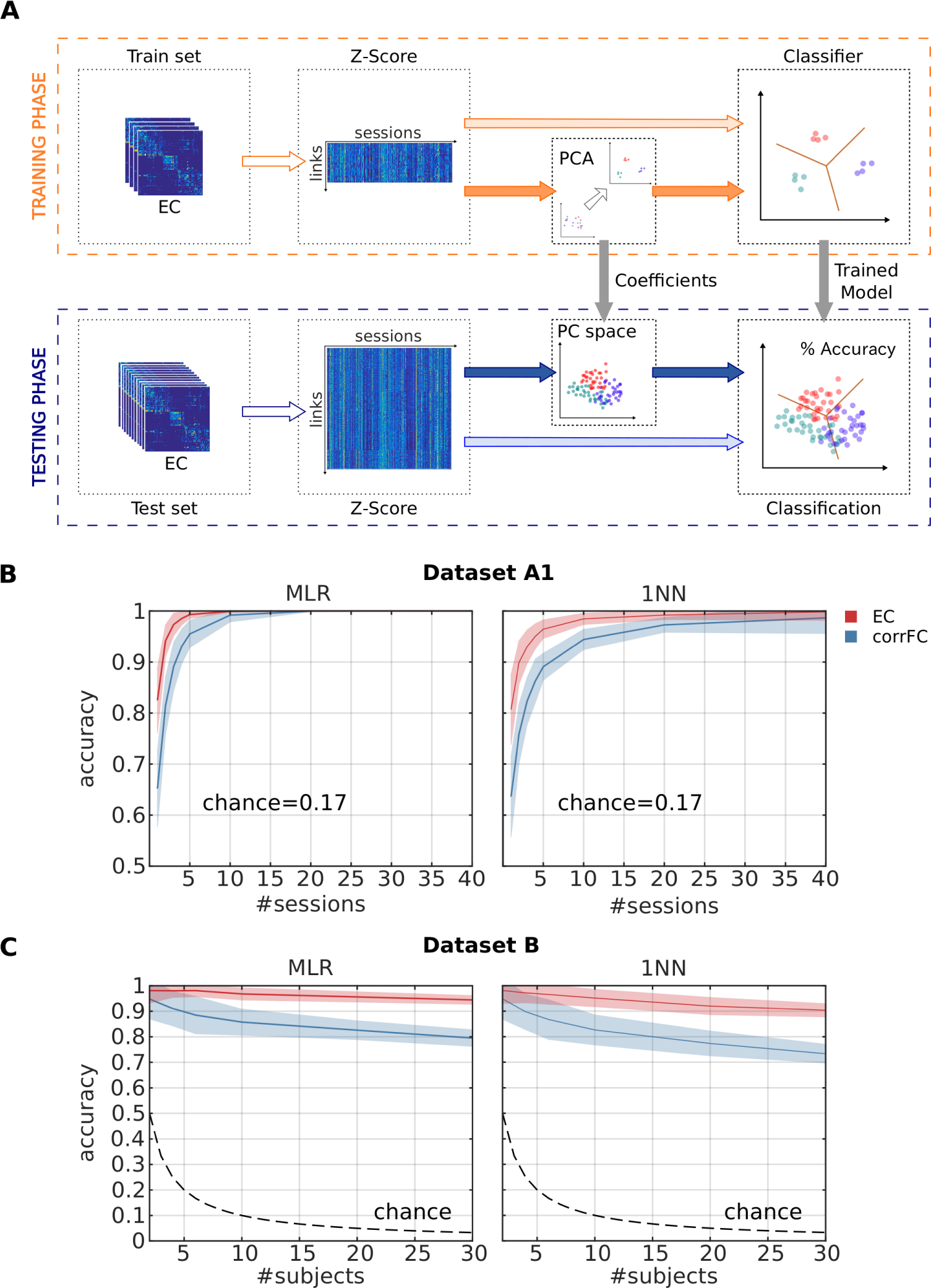
**Comparison between EC and FC in subject identification. A)** Classification pipeline used to assess the generalization of performance. The full set of connectivity measures (here EC) over all fMRI sessions was split into two groups: a train set and a test set. We use z-scores calculated over the elements of each session matrix (see Eq. (16) in Methods). We trained the classifier—with or without previously applying PCA—and evaluated the classification accuracy on the test set. **B**) Performance of multinomial logistic regression (MLR, left panel) and 1-nearest-neighbor (1NN, right panel) classifiers when increasing the number of sessions per subject used as training set with Dataset A1. The mean (solid curve) and standard deviation (colored area) were calculated for 100 repetitions with cross-validation. C) Same as B when varying the number of subjects using Dataset B, using a single training session per subject (leaving 9 sessions per subject as test test).

We first used Dataset A1 and increased the number of training sessions per subject from 1 to 40 to evaluate how many training sessions are necessary for satisfactory accuracy. As shown in Fig. 3B, EC (in red) outperformed corrFC (in blue) by more than one standard deviation (shaded area around the curve), for both MLR and 1NN. For both classifiers, the Mann-Whitney test over 100 repetitions gives a p-value < 10^−11^ until 10 sessions for the superiority of EC. Moreover, almost perfect classification was reached with MLR for only 5 training sessions, whereas 10-15 were necessary for 1NN. This is important when only a few training sessions per subject are available, as expected with clinical applications. Fig. 3C displays the classification accuracy for Dataset B, used to verify the robustness with respect to the number of subjects to be classified. We trained the classifiers with 1 session per subject and evaluated the performance varying the number of subjects from 2 to 30 (test set comprised the remaining 9 sessions per subject). Again, EC is more robust than corrFC: While performance with corrFC rapidly deteriorates as the number of subjects is increased, classification using EC is barely affected by the number of subjects. Again, the Mann-Whitney test over 100 repetitions gives a p-value < 10^−5^ in all cases (more than two subjects). This is our core technical result: EC and MLR largely outperform corrFC and 1NN, respectively. Other connectivity measures such as FC0 showed similar performance to corrFC (not shown).

Unlike with the analysis of the data structure (Fig. 2E), PCA only marginally increased the classification performance here (Supplementary Figure S5). The difference with Fig. 2E may be explained because here PCA can only be applied to the train set. Another possible reason is that the classification accuracy is already very high with 95% for MLR and EC with 1 training session. As further discussed in Supplementary Figures S5 to S8, subject-specific information in conveyed by many PCs, corresponding to a rather large subspace of the EC links. This supports the use of proper machine learning tools to extract this distributed information.

### 2.4 Signature network of links supporting the classification

Another important advantage of the MLR over *k*NN is its efficiency in characterizing the links that contribute to the classification, which can be used to quantitatively study the signatures, as shown in this section. We used recursive feature elimination (RFE, see Methods for details) to rank the links according to their weight in the classification and then to choose the lowest number of links for which the MLR achieved the maximum classification performance. In comparison, the same procedure with *k*NN would require many more computations, recalculating the closest neighbor for all combinations of links (here the number of links is *p* > 1000; see Methods for further discussion). The resulting support network for dataset A1 had 18 links, compared to 44 links for dataset B. In both cases, subject identification using only those links achieved perfect accuracy (90% of all available sessions were used for training and 10% for testing, see Supplementary Figure S9). The two support networks are shown in Fig. 4A in the same matrix: Remarkably, the networks are very sparse and non-uniformly distributed across the whole brain. This is the signature of the most subject-discriminative ROIs: Frontal and cingulate cortices, as well as the temporal and occipital regions, seem to play a major role here. It is worth noting that the EC adjacency matrix is not symmetric, which implies different roles for nodes as receivers (especially frontal ROIs) or senders (cingulate).

**Figure 4:**
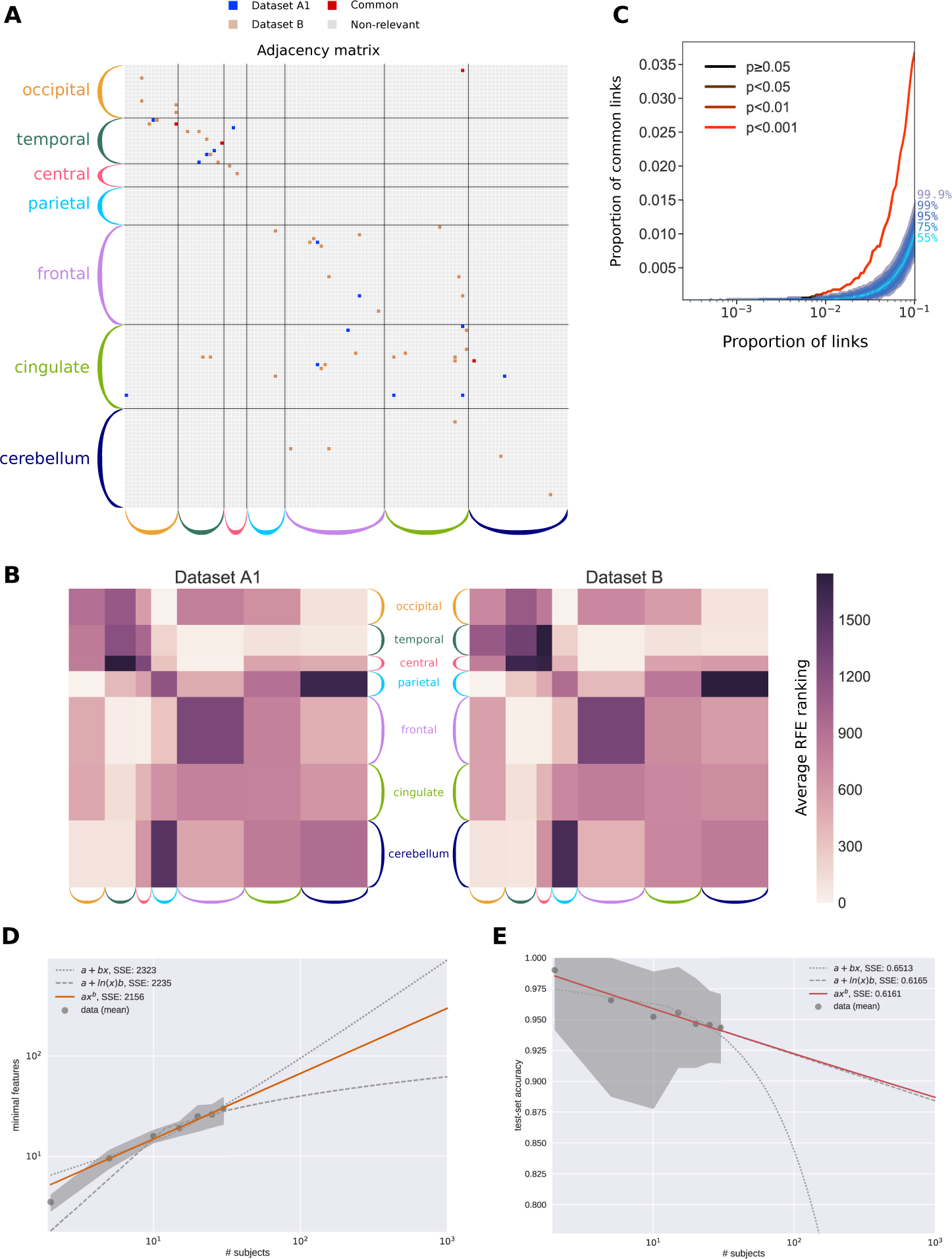
**Support network of EC links to identify subjects and generalization capability. A)** Extracted links that contribute to the classification for Datasets A1 and B, obtained using recursive feature elimination (RFE). The ROIs are grouped in anatomical pools, as detailed in Supplementary Table S1. **B)** Average RFE ranking over the ROI anatomical pools in **A. C)** Overlap between the two signatures for Datasets A1 and B as a function of selected links. The curve represents the amount of common links in the data. Shaded areas represent different quantiles of the surrogate distribution of common links under the null-hypothesis of random rankings. The color of the curve indicates the probability of the corresponding amount of common links under the null-hypothesis (here p-value i 0.001 when considering more than 1% of the total links, namely 40 links). **D)** Extrapolation of the size of the support network in **A** for Dataset B when increasing the number of subjects up to 1000 (x-axis), for 10 repetitions. The shaded area corresponds to 10 repetitions. The curves compare three approximations, the best one displayed in orange (as indicated by the fitting error SSE) corresponds to a sublinear power law (exponent equal 0.6). **E)** Same as **D** for the accuracy based on EC. The plotted data points correspond to EC in Fig. 3C and the variability to 100 repetitions. The best approximation also corresponds to a power law (in orange), which is very close to a lognormal relationship.

The sparsity of the signature in Fig. 4A hides the fact that the rankings for Datasets A1 and B are close (PCC=0.59, p-value ≪ 10^−50^), indicating that similar links convey information about the identity of subjects. Fig. 4B shows the remarkable correspondence of the mean RFE ranking of EC connections for Datasets A1 and B, where the source and target ROIs are grouped anatomically (in color in Fig. 4A). This confirms that the two corresponding signatures coincide at the subnetwork level. Compared to previous studies that based classification on parietal and frontal regions [Finn et al., 2015], we found that temporal and occipital regions are discriminative. An explanation for this difference may lie in that EC consists of directional links and may capture more refined information than FC. To further measure the overlap between these networks, we selected the subset of links with the highest ranking for each dataset and computed the number of common links. Fig. 4C shows that the proportion of common links exceeds by far its expectation under the hypothesis of random rankings (shaded gray area). This indicates a good agreement between the support networks from the two datasets even at the single-link level.

Expecting the size of the support network in Fig. 4A to grow with the number of subjects, we quantified it using Dataset B and found a sublinear relationship, as illustrated in the loglog plot of Fig. 4D. The extrapolation to 100 and 1000 subjects gave network sizes of 53 and 200 links, the latter corresponding to 5% of the total number of EC links (i.e., the signature remained sparse). Likewise, we fitted a curve to predict the EC-based accuracy when increasing the number of subjects (see Fig. 4E), which gave 92% and 88% for 100 and 1000 subjects. These results implied that the complexity of EC biomarkers related to individual variability should not “explode”, which is good news for the generalization capabilities of EC.

### 2.5 Twofold classification of subject identity and behavioral condition

Lastly, we used the knowledge gained from the subject identification to reach our final objective, the extraction of several signatures corresponding to distinct modalities and the evaluation of the overlap between them. From sessions in Dataset C, We extracted a signature for the subject identity and another for the behavioral condition. The twofold classification is schematically depicted in the left panel of Fig. 5A, with three fictive dimensions: The information about subject identity corresponds to the horizontal axis and information about the condition to the vertical axis; The session-to-session variability, which should be ignored, spreads along the depth axis. In this idealized scenario, it is possible to classify a session with respect to both subjects and conditions using different dimensions of the data (middle panel). In the high dimensional case, this occurs when different sets of links support the two classifications. In contrast, the two signatures may involve overlapping subsets of links, corresponding to non-orthogonal axes (right panel) for which the condition “score” for classification (projection on axis) also involves individual information.

**Figure 5:**
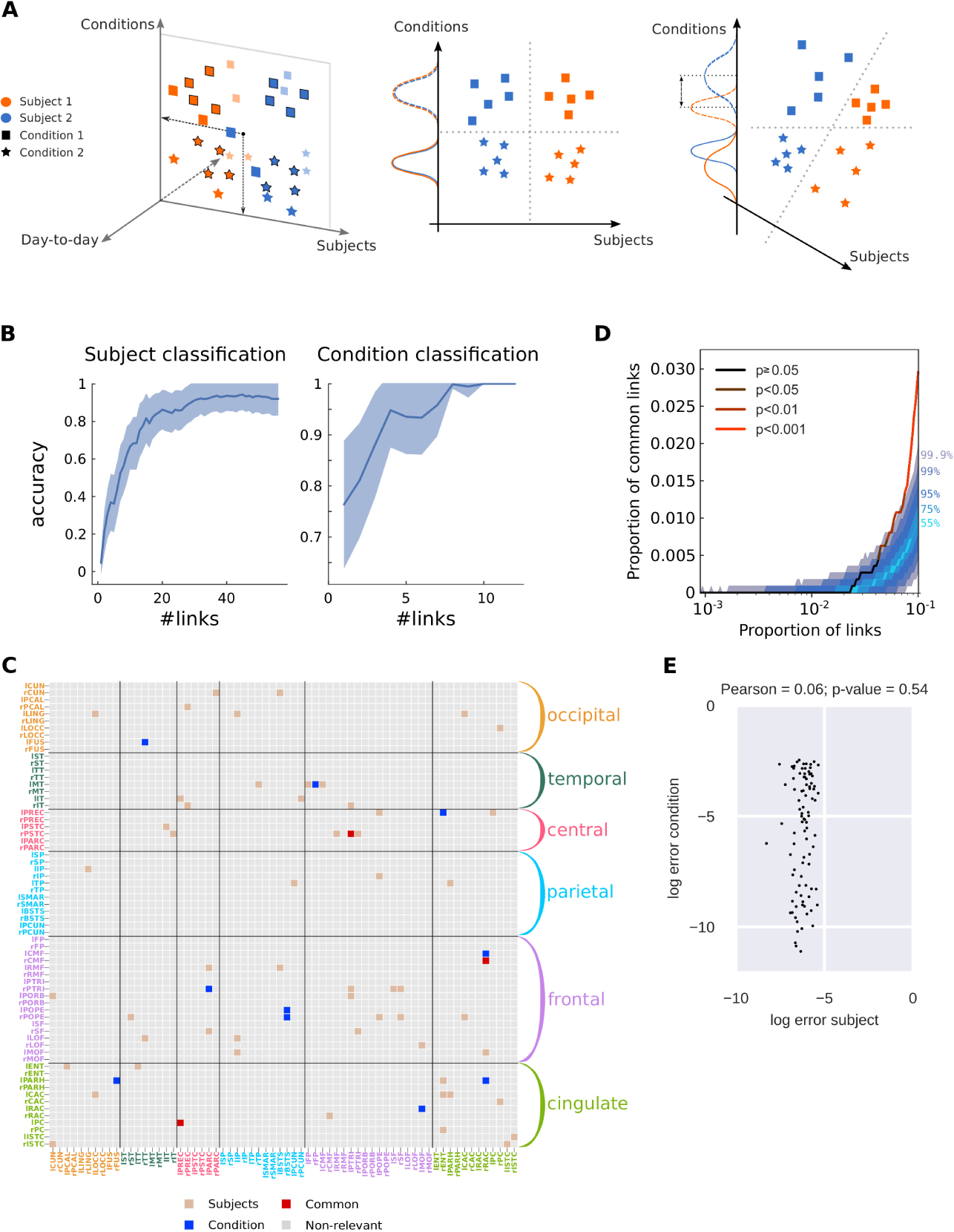
**Twofold discrimination between subjects and conditions using EC. A)** The left panel represents an idealized scheme of the twofold classification where the biomarkers for all sessions (colored symbols) are projected onto two subspaces, one for subjects (orange versus blue) and one for conditions (squares versus stars). The middle and right panels compare the situations where the subspaces (here axes) are orthogonal or not, respectively. In both cases, the dashed separatrices correspond to perfect classification. In the right panel, however, the condition “score” (projected value) mixes information about subjects, as indicated by the non-overlapping solid and dashed distributions. **B)** Performance of the classification for 19 subjects and 2 conditions using Dataset C as a function of number of links. Note the distinct scales for the y-axis, because the subject identification is a harder problem. **C)** Signatures of the most discriminative EC links (estimated with RFE, see text for details) for the twofold classification in B: 54 links for subject classification in brown, 10 for condition classification in blue, 3 common links in red. The ROIs are grouped in anatomical pools, as detailed in Supplementary Table S2. **D)** Proportion of common links between the subject and condition signatures as a function of selected links (in the order of the RFE ranking). Color coding is the same as in Fig. 4B: the two signatures are significantly different, i.e., with a number of common links corresponding to the null hypothesis with p-value ≥ 0.05 (cf. legend) with up to 4% of the total links. **E)** Error of the subject and condition classifiers using the links of the support networks in **C**. The plotted dots represent the 95 sessions. The classifier errors are not correlated, as indicated by the Pearson correlation and corresponding p-value.

MLR with EC achieved very high performance (accuracy ¿ 90%) for subject identification and perfect classification for the condition. We then sought the smallest subsets of links that achieved the maximum performance of each classification using RFE (Fig. 5B), as done before (see Methods for details). Both support networks were again very sparse and distributed across the brain, as can be seen in their adjacency matrix (Fig. 5C). More links are necessary to identify the subjects (57) than the behavioral conditions (13), indicating a higher complexity for the former.

To quantify the overlap between the signatures, we computed the number of common links supporting both subject and condition identifications, when increasing the two subsets according to the order given by RFE as before in Fig. 4C. The overlap fell within the expected values for the null hypothesis up to 30% of all links, as illustrated in Fig. 5D. This indicated that distinct subsets of links were relevant for the subject identities and behavioral conditions, ensuring the “orthogonality” of the two signatures for this dataset. In addition, the few links in common could be removed and replaced by further links in the RFE ranking to further disentangle the signatures. In any case, this suggested a good generalization property of the multivariate signature extraction for several modalities, in the sense that disjoints subsets of links can support the classifications. This application further demonstrated the above mentioned advantage of the MLR classifier equipped with RFE: Working in the original space of links allows us to identify the most discriminative links individually, as well as combinations of them in a subsequent step. In contrast, PCA-like approaches gives PCs corresponding to linear combinations of many links, making it difficult to interpret results at the level of single links.

Another implication of the non-orthogonality of signatures in the right panel in Fig. 5A is that noise in EC estimates may induce correlated errors in the two classifications. To test this idea, we compared the errors of the two classifiers compared to the perfect outcome (probability of 1 for the correct class and 0 for others). We found that the two classifiers for the support networks in Fig. 5C had uncorrelated errors, with a high p-value in Fig. 5E. This further confirmed that the selection of only a few links to support the signature classifiers preserved a twofold classification without bias between the two modalities.

Similar to Datasets A1 and B, subject identification of Dataset C largely concerned the frontal and cingulate systems. Condition identification was also supported by occipital and temporal cortices, which were found to have the strongest activity modulations during movie viewing [Gilson et al., 2017]. We then compared the two corresponding support networks represented in the top panels of Figures 5A and B (the directed nature of links can be appreciated by zooming). Because we only considered two tasks, it was somehow expected that the task network would be much simpler than the support network for the 19 subjects. However, these two support networks also corresponded to distinct types of graphs: The subject network had a large connected component with several central nodes (hubs, indicated by their large size), located in frontal and cingulate regions, in addition to two small components. In comparison, the condition network was segregated into 8 small isolated components. This suggested that elaborate collective dynamics involving high-level ROIs differed across subjects, which might be used as a biomarker to examine its relationship with individual cognitive traits, as was done with FC [Finn et al., 2015].

The bottom plots in Fig. 6 show the lateralization of the support links, stressing the asymmetries between the two hemispheres: Most of the important links are ipsilateral (i.e., within the same hemisphere) and many belong to the left hemisphere for the subject network, whereas they are mainly contralateral for the condition network. This is also in line with strong inter-hemispheric interactions observed during movie viewing for the same dataset, but relying on community analysis [Gilson et al., 2017]. This shows that the non-overlap between the signatures is not only quantitative, but also qualitative in the sense that these signatures involve distinct types of links.

**Figure 6:**
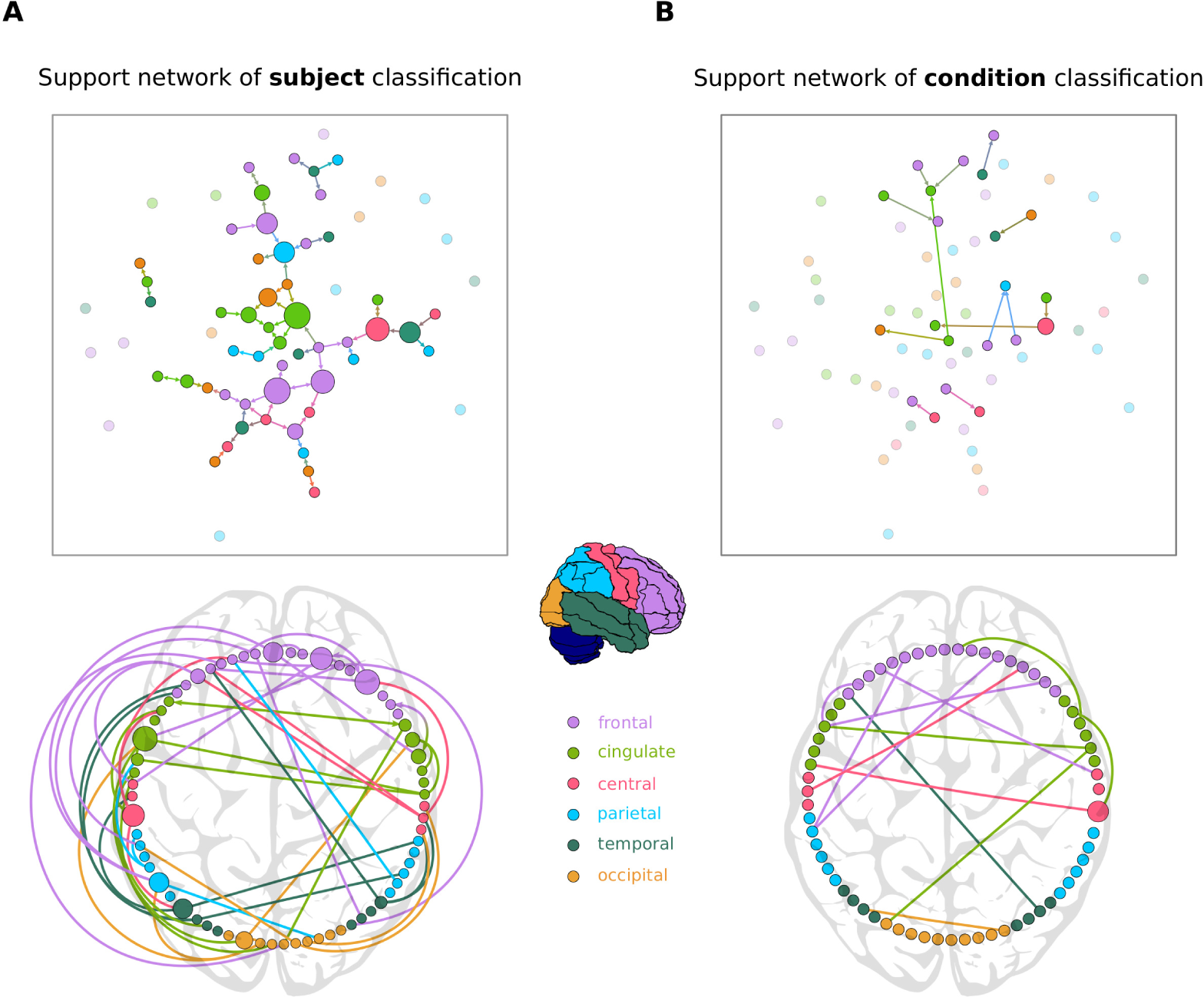
**Support networks of subject and condition classification. A)** The top graph plot represents the 57 most discriminative EC links supporting the classification of subjects (same as in Fig. 3C). The size of each node represents its betweenness centrality in the extracted network. The most central regions are located mainly in the frontal and cingulate cortices. The bottom circular plot shows the asymmetry and lateralization of the network, with more links located in the left hemisphere. Links that are inside the circle correspond to contralateral connections, while links outside the circle correspond to ipsilateral connections. **B)** Similar graph and circular plots as A for the 13 links supporting the classification between the two conditions (resting versus movie viewing). Fewer links are required to reach high accuracy in the condition discrimination: they form a network with many disjoint components and are mainly contralateral, in comparison to the subject classification support network. The color for each link corresponds to its source ROI.

## 3 Discussion

In this study, we have proposed a framework to extract signatures related to various modalities from fMRI data, allowing for a quantification of the interference between signatures. The twofold classification of subjects and behavioral conditions goes beyond previous studies that used FC as a “fingerprint” [Miranda-Dominguez et al., 2014, Finn et al., 2015, Calhoun et al., 2017, Finn et al., 2017]. It is precisely because fMRI signals are dominated to some extent by individual traits [Finn et al., 2015] that a proper multivariate classification is necessary. In particular, we have shown how machine-learning tools such as the MLR enable a quantification of the topological orthogonality between signatures, in addition to efficiently extracting them. Our study also demonstrates that signatures based on EC are very robust to session-to-session variability (much more than with FC) and can be obtained relying on a limited number of sessions, namely 4-5 recording sessions (of 5 minutes each) to classify 40+ other sessions. This corresponds to a train-test ratio of 10-90, which is unusual low; in contrast, most studies use ratio above 50-50, which raises concerns about generalization capabilities [Varoquaux et al., 2017]. Our method yields consistent results for fMRI datasets collected using different scanners and preprocessed with different pipelines, which further supports its general scope. Taken together, our results define a solid ground to scale up the multivariate classification to larger datasets with more subjects and tasks. We now discuss specific points.

We found that very sparse signatures (a few % of all EC links) are sufficient to obtain perfect classification because of the datasets we used, which involved 30 subjects maximum and 2 conditions. The size of the corresponding support networks (Figs. 4A and 5C) is expected to increase with the complexity of the “environment” to be represented, with more subjects and tasks. Open-access resources are becoming available to quantitatively test the framework on larger scales [Zuo et al., 2014, Gordon et al., 2017]. An encouraging result for the generalization capability of EC is the sublinear dependency of the signature size with the subject number (Fig. 4D). The sparsity of signatures means that the complexity of the variability of EC biomarkers remains “controlled” in the sense that it can be described using a number of dimensions much smaller than the numbers of elements in the corresponding category (here subjects or conditions).

The support networks for the twofold classification (subject and condition) show several noticeable differences (Fig. 6). The subject network is large, almost fully connected, distributed over the two hemispheres (with more links within the left one) and concentrated in the cingulate and frontal areas. This suggests subject-specific dynamics between areas involved in high-level functions and overlapping with the default mode network [Raichle et al., 2001]. These discriminative EC patterns may reflect heterogeneities in the interactions between the different neural subsystems (e.g., cingulate to frontal in Fig. 5C) and the propagation of information between them [Ekstrom, 2010, Engel et al., 2013]. As expected with the movie viewing condition studied here, links in the visual and temporal areas are discriminative. In addition, we also found a much higher percentage of contralateral links for condition than subject. This suggests that EC-based biomarkers may also be interpreted in terms of brain communication as was shown recently [Gilson et al., 2017, Senden et al., 2017], beyond simply supporting the classification.

A fundamental advancement of our study is the development of a reliable and well-benchmarked method, extending previous proofs of concept for subject or condition identification [Miranda-Dominguez et al., 2014, Finn et al., 2015, Gonzalez-Castillo et al., 2015]. Our core technical result is that (whole-brain) EC discriminates subjects better than corrFC (Fig. 2 and Fig. 3), which is used in the vast majority of previous studies [Finn et al., 2015, 2017, Kaufmann et al., 2017, Calhoun et al., 2017, Varoquaux et al., 2017, Woo et al., 2017, Xie et al., 2017, Drysdale et al., 2017]. The generalization capability of EC is much more robust than FC when the classification becomes harder (many subjects to identify with few sessions per subject, Fig. 3B-C). This confirms that the BOLD temporal structure—captured by the EC after the bandpass filtering of the BOLD signals—reflects the identity of the subject [Miranda-Dominguez et al., 2014], as previously shown for a task involving attention (or not) [He, 2011] or for wake versus sleep [Mitra et al., 2015]. The use of z-scores in the classification shows that the EC ranking (i.e., which brain connections have large weights among all) conveys the relevant information. This is also in line with our previous studies [Gilson et al., 2017, Senden et al., 2017] that showed how changes in task-evoked fMRI activity is captured by whole-brain EC. Moreover, the directed nature of EC reflects the propagation of BOLD signals and can be interpreted in terms of brain communication. Our results thus support a change in biomarkers used for multivariate classification, where EC unfolds the relevant information of BOLD signals in a suitable space. Here the focus was on EC because it performed better than Σ estimates for the resting-state fMRI, but it has been shown that Σ may be strongly affected by tasks [Gilson et al., 2017], so Σ might further improve the classification for conditions, reflecting sensory stimuli.

On the machine learning side, our results show that the MLR classifier performs better than 1NN (Fig. 3B-C), as well as *k*NN with *k* ≥ 2 (results not shown). This suggests that the EC/FC pools related to subjects/conditions are linearly separable in their high dimensional spaces, in a even easier manner for EC (related to the better performance). Unlike the *k*NN classifier [Finn et al., 2015], the MLR discards uninformative links with little reliability to discriminate the desired modalities, so taking whole-brain EC with many dimensions (links) without a-priori selection is not an issue. In addition, we have shown how RFE can be used to to quantitatively study the signatures and assess the orthogonality between them as it provides access to the link-level contributions to classification (Fig. 5D-E). Therefore, the multivariate classification can be implemented using distinct subsets of links, discarding those that mix several modalities. Together, these results underline the importance of adequate machine learning tools to obtain a powerful and flexible framework that can scale up.

Formally, our whole-brain dynamic model is a continuous-time network with linear feedback that incorporates topological constraints from SC. EC is estimated using a gradient descent (or Lyapunov optimization) that takes into account the network feedback and can be very efficiently calculated for the whole brain with 100 ROIs and each session with 300 time points per ROI [Gilson et al., 2016, 2017]. Each session gives a parameter estimate (for each EC link), whose distributions across subjects and conditions are used for classification. Our dynamic model and estimation procedure are simpler than the dynamic causal model with hemodynamics and Bayesian machinery [Friston et al., 2003, Stephan et al., 2004], which has been used for classification relying on a few ROIs only [Brodersen et al., 2011]. Nonetheless, they provide powerful signatures in a much richer (high-dimensional) space that can be used for modality discrimination. Our study focused on two coarse parcellations covering the whole brain [Tzourio-Mazoyer et al., 2002] or cortex [Hagmann et al., 2008]. Although the two parcellations were applied to different datasets, we did not observe significant difference in the performance of the classifiers. Much work has been done recently to correct the bias due to the use of specific parcellations [Da Mota et al., 2014]; for our purpose, more refined parcellations may entail better discriminability in higher-dimensional spaces, but raise issues for the EC estimation robustness. This motivated us to choose rather coarse parcellations with large ROIs as a first step. We have used a generic SC with 30% density, ensuring a sufficiently rich potential biomarker with thousands of estimated EC links in total. Inaccuracies about SC may affect some % of all links, but this is unlikely to hardly affect the collective predictive power of EC. Moreover, preprocessing using PCA was surprisingly found to have very little influence on the performance. Nonetheless, PCA may be useful for datasets with larger number of subjects and conditions [Preti et al., 2017]. We also tried the classification procedure with additional global signal regression of the BOLD signals and results were similar in terms of performance. Although refining the preprocessing pipeline may improve the (already excellent) classification performance, we expect EC to perform better in general.

Applications of neuroimaging to characterize brain disorders at the patient level are emerging [Matthews and Hampshire, 2016, Yahata et al., 2017]. The development of personalized medicine with tailored therapeutic protocols [Shen, 2014]—to optimize recovery and minimize adverse effects—requires quantitative tools that deliver a precise diagnostic of the patient’s evolution. Our proposed scheme is to follow a patient’s trace over time in the (high-dimensional) EC space: Extending the diagram in Fig. 5A, the classification should involve a 4-dimensional space (session-to-session variability to discard, subject, task and pathology), the latter dimension corresponding to healthy versus pathological states. Here the goal of the multivariate signatures is to ensure that the pathology one is not “polluted” by other modalities. One (or several) pathology-specific signature(s) would be extracted from resting-state [Greicius, 2008] or task-evoked fMRI, as specific tasks may indeed reveal powerful signatures for certain pathologies, e.g., memory exercises for Alzheimer [Kurth et al., 2015]. We expect our framework to bridge the gap between the two types of recent approaches that dealt with either side of individual traits, but not both at the same time: 1) A recent prospective study on the evolution of psychiatric disorders emphasized individual specificities in the FC stabilization during childhood (but irrespective of the disease) [Kaufmann et al., 2017]; 2) Group-averaging is often used to ignore the individual differences and obtain pathology-specific signatures [Drysdale et al., 2017]. The generalization capability of prediction methods to future (unseen) data [Hughes, 1968, Calhoun et al., 2017, Varoquaux et al., 2017, Woo et al., 2017] is crucial in this clinical context. To this end, our method appear suitable for disentangling diverse signatures, while properly conditioning out the day-to-day fMRI variability (as uninformative intrinsic noise). This provides a practical solution to the recent criticism that “a major reason for disappointing progress of psychiatric diagnostics and nosology is the lack of tests which enable mechanistic inference on disease processes within individual patients” [Stephan and Mathys, 2014].

## 4 Methods and Materials

### 4.1 Description of fMRI datasets

Three datasets acquired at different locations were used in this work:

- Dataset A was acquired for the Day2day project [Filevich et al., 2017] at the Max Planck Institute for Human Development (Berlin, Germany) of 40-50 resting-state sessions recorded from the 6 subjects over 6 months. The uniqueness of this data lies in the capability to have statistically valid evaluation of the session-to-session variability for single subjects.
- Dataset B is publicly available and is part of the Consortium for Reliability and Reproducibility (CoRR) [Zuo et al., 2014]. We used this dataset to generalize the results of the discriminability when increasing the number of subjects (up to 30 subjects).
- Dataset C [Mantini et al., 2012] was recently analyzed using our model-brain dynamic model to extract effective connectivity [Gilson et al., 2017]. We used it to perform a twofold classification with respect to both subjects and conditions (resting-state versus movie viewing).

In this section, we provide details about the acquisition of the blood-oxygen-level dependent (BOLD) signals for the three datasets.

#### 4.1.1 Dataset A

This dataset has two parts. The first part (A1) is longitudinal and consists of resting-state fMRI sessions from 8 subjects (age 24-32, 6 female). 2 subjects (one male, one female) were not able to continue the study and were discarded. The other 6 subjects underwent scanning between 40 and 50 times along a period of 6 months. The second part of the dataset (A2) was acquired during the same period of time. A total 50 subjects (age 18-32, all female) were scanned during a single fMRI session each using the same MRI sequences. Participants were free of psychiatric disorder according to a personal interview—mini-international neuropsychiatric interview [Sheehan et al., 1998]—and had never suffered from a mental disease. The study was approved by the local ethics committee (Charite University Clinic, Berlin). Participants were instructed to remain with their eyes closed and data acquisition had to be constrained to 5 min per scan due to experimental limitations.

Images were acquired on a 3 T Magnetom Trio MRI scanner system (Siemens Medical Systems, Erlangen, Germany) using a 12-channel radiofrequency head coil. Functional images were collected using a T2*-weighted echo planar imaging (EPI) sequence sensitive to BOLD contrast (TR = 2000 ms, TE = 30 ms, image matrix = 64 × 64, FOV = 216 × 216 × 129 mm^3^, flip angle = 80°, bandwidth=2042 Hz/pixel, voxel size=3 × 3 × 3 mm^3^, 36 axial slices using GRAPPA acceleration factor. Structural images were obtained using a three-dimensional T1-weighted magnetization-prepared gradient-echo sequence (MPRAGE) based on the ADNI protocol (www.adni-info.org): repetition time (TR) = 2500 ms; echo time (TE) = 4.77 ms; TI = 1100 ms, acquisition matrix = 256 × 256 × 192 mm^3^, flip angle = 7°; bandwidth=140 Hz/pixel, 1 × 1 × 1 mm^3^ voxel size.

##### Pre-processing

The data was preprocessed using SPM5 (Wellcome Department of Cognitive Neurology, London, UK) and DPARSF/DPABI [Yan et al., 2016] after discarding the first 10 volumes of each session. It included: slice timing and head-motion correction (6 parameters spatial transformation), spatial normalization to the Montreal Neurological Institute (MNI) template, and spatial filtering of 4 mm FWHM. Linear trends were removed from the fMRI time courses before band-pass filtering (0.01-0.08 Hz). The data was parcellated using the automated anatomical labeling (AAL) atlas [Tzourio-Mazoyer et al., 2002] into 116 regions of interest (ROIs), which includes the whole cortex and the cerebellum.

#### 4.1.2 Dataset B

This dataset consists of 10 fMRI resting-state sessions acquired from 30 healthy participants every three days for one month [Zuo et al., 2014]. Each session lasted 10 minutes. To minimize head movement, straps and foam pads were used to fix the head snugly during each scan. The participants were instructed to relax and remain still with their eyes open, not to fall asleep, and not to think about anything in particular. The screen presented a fixation point and after the scans, all the participants were interviewed, and none of them reported to have fallen asleep in the scanner. The time of day of MRI acquisition was controlled within participants.

Recording sessions were performed using a GE MR750 3.0 Tesla scanner (GE Medical Systems, Waukesha, WI) at CCBD, Hangzhou Normal University. T2-weighted echo-planar imaging (EPI) sequence was performed to obtain resting state fMRI images for 10 minutes using the following setup: TR = 2000 ms, TE = 30 ms, flip angle = 90°, field of view = 220 × 220 mm^2^, matrix = 64 × 64, voxel size = 3.4 × 3.4 × 3.4 mm^3^, 43 slices. A T1-weighted fast spoiled gradient echo (FSPGR) was used with the following protocol: TR = 8.1 ms, TE = 3.1 ms, TI = 450 ms, flip angle = 8°, field of view = 256 × 256 mm^2^, matrix = 256 × 256, voxel size =1 × 1 × 1 mm^3^, 176 sagittal slices) was carried out to acquire a high-resolution anatomical image of the brain structure.

##### Pre-processing

Dataset B was preprocessed with SPM12 (Wellcome Trust Centre for Neuroimaging, London, UK) and DPARSF/DPABI [Yan et al., 2016]. The first 5 fMRI volumes were discarded in order to let the BOLD signal reach stability. The pre-processing pipeline included: slicetiming correction, realignment for motion correction, co-registration of the T1 anatomical image to the mean functional image, detrending, regression of 6 movement parameters, 5 principal component analysis (PCA) white matter and CSF Compcorr, and spatial normalization to MNI coordinates. Scrubbing with Power 0.5 and linear interpolation was applied. This data was also parcellated into 116 ROIs using the AAL parcellation [Tzourio-Mazoyer et al., 2002] and time courses were band-pass filtered between 0.01 and 0.08 Hz, as done with Dataset A. No further global signal regression and spatial smoothing were applied.

#### 4.1.3 Dataset C

We used a third dataset to study the discrimination between subjects and conditions. In this case, a total of 22 subjects (age 20-31, 15 females) were scanned during rest with eyes opened and natural viewing condition. Volunteers were informed about the experimental procedures, which were approved by the Ethics Committee of the Chieti University, and signed a written informed consent. In the resting state, participants fixated a red target with a diameter of 0.3 visual degrees on a black screen. In the natural viewing condition, subjects watched and listened to 30 minutes of the movie ‘The Good, the Bad and the Ugly’ in a window of 24 × 10.2 visual degrees. Visual stimuli were projected on a translucent screen using an LCD projector, and viewed by the participants through a mirror tilted by 45 degrees. Auditory stimuli were delivered using MR-compatible headphones. For each subject, 2 and 3 scanning runs of 10 minutes duration were acquired for resting state and natural viewing, respectively.

The BOLD signals were acquired with a 3T MR scanner (Achieva; Philips Medical Systems, Best, The Netherlands) at the Institute for Advanced Biomedical Technologies in Chieti, Italy. The functional images were acquired using T2*-weighted echo-planar images (EPI) with BOLD contrast using SENSE imaging. EPIs comprised of 32 axial slices acquired in ascending order and covering the entire brain with the following protocol: TR = 2000 ms, TE = 3.5 ms, flip angle = 90°, in-plane matrix = 230 × 230, voxel size = 2.875 × 2.875 × 3.5 mm^3^. For each subject, 2 scanning sessions of 10 minutes duration were acquired for resting state and 3 for natural viewing. Each run included 5 dummy volumes, allowing the MRI signal to reach steady state and the subsequent 300 functional volumes were used for the analysis. Eye position was monitored during scanning using a pupil-corneal reflection system at 120 Hz (Iscan, Burlington, MA, USA). A three-dimensional high-resolution T1-weighted image was acquired for anatomical referencing using an MPRAGE sequence with TR = 8.1 ms, TE = 3.7 ms, voxel size=0.938 × 0.938 × 1 mm^3^ at the end of the scanning session.

##### Pre-processing

Data were preprocessed using SPM8 (Wellcome Department of Cognitive Neurology, London, UK), including slice-timing and head-motion correction (see Methods in Gilson et al. [2017] for details), co-registration between anatomical and mean functional image, and spatial normalization to MNI stereotaxic space (Montreal Neurological Institute, MNI) with a voxel size of 3 × 3 × 3 mm^3^. Mean BOLD time series were extracted from N = 66 regions of interest (ROIs) of the brain atlas used in [Hagmann et al., 2008] for each recording session.

The data are available at github.com/MatthieuGilson/EC_estimation. The discarded subjects in the present study are 1, 11 and 19, among the 22 subjects available online (numbered from 0 to 21). The same subjects were discarded in our recent study [Gilson et al., 2017] because of abnormally high BOLD variance.

#### 4.1.4 Structural connectivity (SC)

For all models, we used a generic matrix of structural connectivity to determine the skeleton of the effective connectivity (i.e., existing connections). For For Dataset C, structural connectivity for Dataset C was estimated from diffusion spectrum imaging (DSI) data collected in five healthy right-handed male participants [Hagmann et al., 2008]. The gray matter was first parcellated into the *N* = 66 ROIs, using the same low-resolution atlas used for the FC analysis. For each subject, we performed white matter tractography between pairs of cortical areas to estimate a neuro-anatomical connectivity matrix. In our method, the DSI values are only used to determine the skeleton: a binary matrix of structural connectivity (SC) obtained by averaging the matrices over subjects and applying a threshold for the existence of connections. The strengths of individual intracortical connections do not come from DSI values, but are optimized as explained below. For Datasets A and B, the generic SC corresponded to the AAL parcellation with *N* = 116 ROIs [Tzourio-Mazoyer et al., 2002]. A similar pipeline was used with diffusion tensor imaging.

It is known that both tractography and DTI underestimate inter-hemispheric connections [Hagmann et al., 2008]. Homotopic connections between mirrored left and right ROIs are important in order to model whole-cortex BOLD activity [Messe et al., 2014]. For both SC matrices, we added all possible homotopic connections, which are tuned during the optimization as other existing connections.

### 4.2 Connectivity measures and model estimates

Here we provide details about the calculation of the functional and effective connectivity measures introduced in Fig. 1.

#### 4.2.1 Empirical measures of functional connectivity

For each fMRI session, the BOLD time series is denoted by 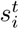 for each region 1 ≤ *i ≤ N* with time indexed by 1 ≤ *t* ≤ *T* (time points separated by a TR=2 seconds). The time series were first centered by removing—for each individual ROI *i*—the session mean 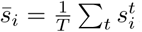. Following [Gilson et al., 2016], the spatiotemporal FC corresponds to covariances calculated as:

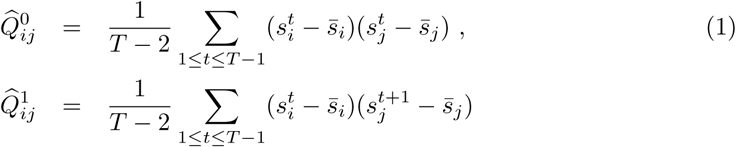

The classical BOLD correlations (corrFC in the main text) correspond to

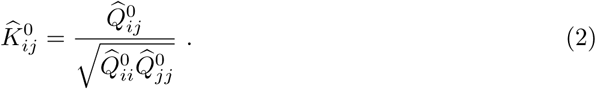

#### 4.2.2 Model of cortical dynamics

The model uses two sets of parameters to generate the spatiotemporal FC:

- The network effective connectivity (EC) between the ROIs (cf. Fig. 2 in the main text) is denoted by the matrix *C* in the following equations. Its skeleton is determined by the SC matrix, but not its weight values: Weights for absent connections are kept equal to 0 at all times, but weights for existing connections are estimated from FC matrices for each session.
- The local variability is described by the variances (1 per ROI) on the diagonal of the matrix Σ.

The model FC comes from the propagation of the local variability—inputed to every ROI—that propagates via EC, generating network feedback.

Formally, the network dynamics is described by a multivariate Ornstein-Uhlenbeck process, where the activity variable *x_i_* of node *i* decays exponentially with time constant *τ_x_*—estimated using Eq. (7)—and evolves depending on the activity of other populations:

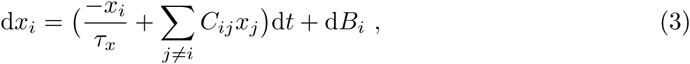

where d*B*_*i*_ is equivalent to white noise with covariance matrix Σ (formally a Wiener process); note that only variances on the diagonal are non zero here.

The simplicity of the model allows for an analytic (feedforward) estimation of the covariances 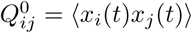 and 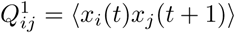, which must reproduce the empirical 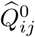 and 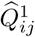, respectively. In practice, we use the two time shifts 0 and 1 TR, because this gives sufficient information to uniquely infer the network parameters (in the theory). Assuming known network parameters *C* and Σ, the matrix *Q*^0^ can be calculated by solving the Lyapunov equation (for example using the Bartell-Stewart algorithm):

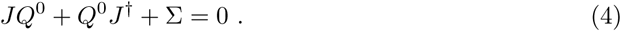

For *Q*^1^, it is simply given by

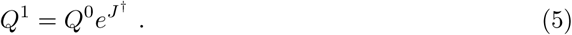

Here *J* is the Jacobian of the dynamical system and depends on the time constant *τ_x_* and the network effective connectivity:

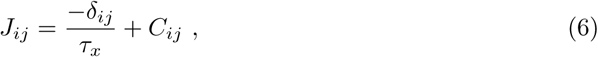

where *δ_ij_* is the Kronecker delta and the superscript † denotes the matrix transpose; note also that *e*^*J*†^ is a matrix exponential.

#### 4.2.3 Parameter estimation procedure

Here we provide details about the Lyapunov optimization (or gradient descent) which is used to tune the model FC to the empirical FC of a given fMRI session. Although it differs from a maximum-likelihood estimate as classically used for multivariate autoregressive processes, it provides a single estimated value for each model parameter.

For each individual and session, we calculate the time constant *τ_x_* associated with the exponential decay of the autocovariance averaged over all ROIs:

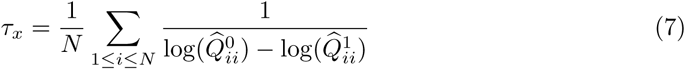

This is used to “calibrate” the model, before its optimization.

In order to invert the model (i.e., for the model FC to reproduce the experimental FC), we iteratively tune the parameters to reduce the model error defined as

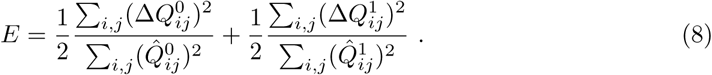

Here each term—for FC0 and FC1—is the matrix distance between the model and the data observables, normalized by the norm of the latter; for compactness, we have defined the difference matrices are 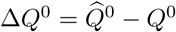 and 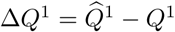.

The idea behind the tuning algorithm is to start from zero connectivity *C* = 0 and homogeneous Σ, then calculate the model *Q*^0^ and *Q*^1^ using the desired Jacobian update given by

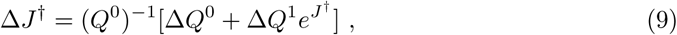

which decreases the model error *E* at each optimization step, similar to a gradient descent. The best fit corresponds to the minimum of *E*. Finally, the connectivity update is

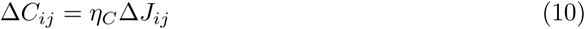

for existing connections only; other weights are forced at 0. We also impose non-negativity for the EC values during the optimization. To take properly the effect of cross-correlated inputs into account, we use the Σ update as in Gilson et al. [2017]:

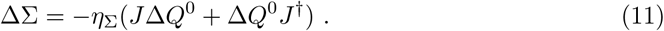

As with *C* for non-existing connections, off-diagonal elements of Σ are kept equal to 0 at all times.

In numerical simulations, we use *η_C_* = 0.0005 and *η*_Σ_ = 0.05. Further details about the derivation of the optimization updates are provided in Gilson et al. [2016].

The optimization code is available with Dataset *C* at github.com/MatthieuGilson/EC_estimation.

#### 4.2.4 Comparison of the model to state-of-the-art dynamic models to interpret fMRI data

Compared to dynamic causal modeling [Friston et al., 2003, Friston, 2011], our model makes the simpler assumption of linearity for the local dynamics. Doing so, it ignores an explicit modeling of the mapping between the neuronal activity and the BOLD signals [Stephan et al., 2004]. Moreover, it uses a simple model of local variability (Wiener process) related to Σ to generate FC than recent development of DCM for resting state [Friston et al., 2014]. In exchange for this simplicity, we obtain a very efficient estimation procedure for networks of about 100 ROIs with 30% density, yielding ∼ 3000 EC parameters. It is also worth noting that the objective function for our framework are the BOLD covariances, which are canonically related to the BOLD cross-spectrum used in the DCM for resting state [Friston et al., 2014].

### 4.3 Analysis and classification of vectorized EC and corrFC

In this section, we provide details about the analysis of the corrFC and estimated EC matrices that are compared across sessions for subject and condition identification. As illustrated in Fig. 1C, the connectivity measure for each session *k* was transformed into a vector *v^k^* by extracting the lower triangle for corrFC, and by applying the SC mask for EC. Following the literature in machine learning, we refer to a connectivity measure for a single session as a sample. The size of the samples *v^k^* is *p* = 6670 for an FC session and *p* = 4056 for an EC session with Datasets A and B (corresponding to *N* =116 ROIs). For Dataset *C*, we used only EC with *p* = 1114 vector elements (for *N* = 66). *Note that we use a slightly different indexing in this section compared to the previous one: In the following i refers to a link in the vector* 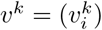, *similarly to the pair* (*i,j*) *for a matrix element C_ij_ before*.

#### 4.3.1 Similarity between sessions

We used Pearson correlation coefficient (PCC) as a measure of similarity, both within and between subjects. For a pair of sessions *k* and *l* as reported in Fig. 2A and B, this is:

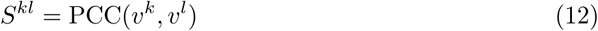

The distribution of within-subject similarity (WSS) in Fig. 2B was obtained by using all pairs of vectors *v^k^* and *v^l^* with *k* ≠ *l* from the same subject, in both Dataset A1 and B. To compute the distribution of between-subject similarity (BSS), all possible combinations of vector pairs *v^k^* and *v^l^* from distinct subjects were used.

#### 4.3.2 Dimensionality analysis

To study visually how the variability of the data is spread over in the space with high dimension p, we applied principal component analysis (PCA) to extract the main dimensions (principal components, or PCs) that capture the largest portion of the data variance. In Fig. 2C that compares corrFC and EC, PCA was applied to the whole Dataset A1 (6 subjects, 40-50 sessions per subject) and the first 6 PCs were retained. Each panel corresponds to PC1 to PC3 on the one hand, and PC4 to PC6 on the other hand.

#### 4.3.3 Silhouette coefficient

The silhouette coefficient [Rousseeuw, 1987] is defined for each vectors *v^k,s^* with indices *k* for the session and *s* for the subject. Here, each subject *s* is a cluster and the similarity in Eq. (12) is taken as the metric, but the indices *k* and *l* are replaced by doublets of the type (*k, s*) here. For a given sample *k, s*, we have the average similarity within his own cluster *s* defined as

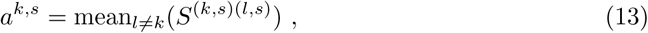

and the maximum—over all other clusters—of the same average similarity, but with elements from another cluster:

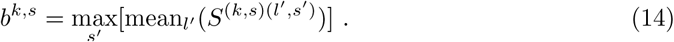

The silhouette is then given by the following contrast between the cohesion of the element within its cluster (*a^k,s^*) and the separation from other clusters (*b^k,s^*):

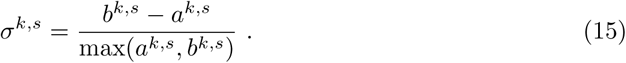

Values of silhouette range to 1 for fully separated clusters to -1 for fully overlapping clusters. These values correspond to the violin plots without PCA in Fig. 2E.

In Fig. 2D and E, the silhouette coefficient were computed for each point in these clouds in a 6-dimensional PC space for Dataset A1. The reason for choosing the first 6 PCs is because the mean silhouette coefficient of EC data reaches a maximum, before decreasing (Supplementary Figure S3, left panel). The same method was applied to the Dataset B (30 subjects, 10 sessions per subject), for which the retained maximum was 30 PCs (Supplementary Figure S3, right panel).

#### 4.3.4 Within-session z-scoring

For the classification in Figures 3 and 4, the values of the EC and corrFC links were z-scored within each session, using the mean and standard deviation of the corresponding vectorized connectivity measure *v^k^* in Fig. 1C:

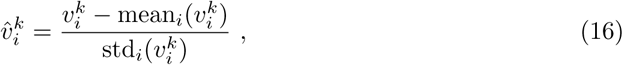

where the vector elements are indexed by *i*, corresponding to a link in the EC or corrFC matrix. The z-scored vectors 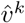 are the inputs of the classifiers, which means that the classification relies on the ranking of the vectorized elements *v^k^* rather than the absolute values of their elements. This is important to understand our claim in the Discussion about where the discriminative information is: The repartition of the weak/strong EC weights across the brain is different across subjects, which is picked up by the algorithms.

#### 4.3.5 Classification of sessions to attribute them to subjects

Fig. 3A shows the classification procedure applied to identify subjects using connectivity measures and estimates. First, a fixed number of sessions per subject were selected from each data set, corrFC and EC matrices. These matrices were vectorized (recall Fig. 1C) and individually z-scored using Eq. (16), using the mean and standard deviation of each *v^k^*. Then, the corresponding classifier—1-nearest-neighbor (1NN) or multilinear logistic regression (MLR)—was trained using two different approaches. In the main text, results are presented without applying PCA as a preprocessing step. This is motivated because PCA does not significantly improve the classification performance for the MLR. Results about classifier with PCA are discussed in Supplementary Figures S5 to S8.

In all cases, the accuracy of a classifier is evaluated by its prediction of samples in the test set, as a cross-validation. We used Dataset A1 to study the effect of increasing the samples in the train set, and Dataset B for increasing the number of subjects. The curves in Fig. 3B and C were obtained after iterating over different sessions and subjects 100 times with this cross-validation (mean and standard deviation are plotted). The same method was also applied to Dataset C by training two MLR classifiers, one for subjects and one for conditions. Both classifiers are available in the scikit-learn package (http://scikit-learn.org, python language).

##### INN classifier

A *k*NN classifier is a technique that assigns to a new sample the class to which belong the majority of its *k* closest neighbors. In our case, we use *k* = 1 with a single nearest neighbor. Moreover, we use the PCC-based similarity measure in Eq. (12) as the metric to evaluate the inverse distance between two samples (here sessions). Like with clustering algorithm in general, closest samples (i.e., most similar sessions) are grouped together. Because the PCC similarity is not linear, it can be considered as a non-linear classifier, in comparison to a linear classifier such as a perceptron or a MLR. In practice, the database has an equal number of sessions per subject—either vectorized and z-scored corrFC or EC—ranging from 1 to 40 for Dataset A1 in Fig. 2B. The identity of the each target session *k* from the test set is predicted by the identity of the most similar session from the test set (*S*^*k*,1^, *S*^*k*,2^, …, *S^k,D^*) with D the size of the database, as illustrated in Supplementary Figure S4 for 1 session per subject as database (and corresponding to the results presented in Fig. 3B and C).

##### MLR classifier

The MLR classifier is a classical tool in machine learning. The parameters (or regressors) of the model are adjusted in order to predict the probabilities of new samples of belonging to each category or class. It relies on the following logistic function that relates to the probability for the vectorized connectivity measure *v^k^* (with elements indexed by *i*) for session k to be in the subject class *s*:

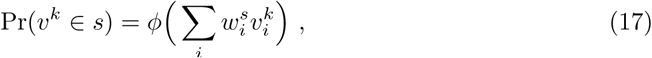

where *ϕ* is the sigmoid function (ranging from 0 to 1). The training is performed by a regression to find the classification weight 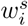 such that Pr(*v^k^* ∈ *s*) discriminates the class *s* against the last subject *s*′. A refinement that does not appear in Eq. (17) is that there is an extra weight to correct for the possibly non-zero mean of the samples *v^k^*. Note also that for there are *M* – 1 regressors for *M* subjects, such that the weights are well constrained. In practice, we used train sets with equal numbers of sessions per subject.

##### Preprocessing using PCA

PCA is a preprocessing step commonly used in machine learning to remove noise while keeping the dimensions that capture most of the variability of the data. This implies that the largest part of the data variances captures the relevant information for the classification. After applying PCA, the original high-dimensional z-scored vectors 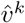 from Eq. (16) are projected into a space of lower dimension determined by a number of PCs. The performance of the classification with PCA thus increases with the number of PCs until saturation (Figure S6), which indicates the point when subsequent PCs contain redundant or irrelevant information for the classification.

##### Extraction of discriminative support networks

We examined which links strongly contributed to the classification, in a similar fashion to PCs as mentioned above. The motivation was that individual links might be mixed in PCs and appear redundantly in several PCs that significantly contributed to the classification (Figures S7 and S8). Therefore, we evaluated the contribution of individual links that supported the classification, forming networks of most discriminative elements in EC (as represented in Figures 3D and 4C). In order to extract these support networks, we employed a commonly used method in machine learning, recursive feature elimination (RFE), to rank the links—taken as features—according to their relevance for the classification [Guyon et al., 2002].

For each application of the RFE algorithm, the train set was composed of 90% randomly chosen samples (to capture the full variability of the data) and test set of the 10% remaining samples. After fitting the MLR classifier, RFE removes the link with the smallest classification weight 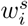 in the MLR formula in Eq. (17), which measures the contribution of the link to the classification. The removing procedure is repeated recursively on the shrinking subset of links until only one is left. This gives a ranking for the links according to their relevance for the classification. We then evaluated the accuracy of MLR on the test set when increasing the number of features following the order given by the RFE ranking. This training and testing procedure was repeated 100 times with different train and test sets each time. We selected the number of features for which the mean test set accuracy was maximum. In order to find the maximum we chose the number of features for which the numerical derivative of the mean was less than 10^−6^. In order to reduce the impact of fluctuations due to the random selection of samples, we smoothed the curve of means with a rolling average of width 2 features. Since the accuracy is expected to increase initially as a function of the number of features and then either saturate or decrease, this method allows for finding the number of features for which the classifier performance is maximum or adding more features has no practical benefit.

In contrast, *k*NN cannot be easily used for RFE since it does not estimate weights associated to links. Therefore, for *k*NN to be used with RFE, one needs to put the model in a wrapper to compare the effect of removing each combination of links on the performance. However, given the high amount of features of our setting, wrappers cannot be evaluated on all subsets of features (∼ 10^300^ tests would be required for 1000 features). Wrappers may rely on approximate greedy algorithms, for example, eliminating the feature that scores worst. It is well known that greedy algorithms might produce inappropriate solutions if the problem does not have optimal substructure. In addition the computation time almost scales as *p*^2^ with *p* the number of links, while for RFE it is linear in *p*.

#### 4.3.6 Software tools

The computer code for the model optimization and classification is available online with Dataset C at github.com/MatthieuGilson/WBLEC_toolbox. It is written in the open-source language python and uses the numpy and scipy libraries, as well as scikit-learn library for machine-learning routines (http://scikit-learn.org).

Connectivity measures and estimates, as well as similarity analyses were performed in MATLAB 2016 (TM). Network plots in Fig. 6 of the main text were done in Gephi 0.9.1 (http://gephi.org).

## 5 Acknowledgments

## References

K. Amunts, M. J. Hawrylycz, D. C. Van Essen, J. D. Van Horn, N. Harel, J.-B. Poline, F. De Martino, J. G. Bjaalie, G. Dehaene-Lambertz, S. Dehaene, P. Valdes-Sosa, B. Thirion, K. Zilles, S. L. Hill, M. B. Abrams, P. A. Tass, W. Vanduffel, A. C. Evans, and S. B. Eickhoff. Interoperable atlases of the human brain. Neuroimage, 99:525–532, Oct 2014. doi: 10.1016/j.neuroimage.2014.06.010.

A. J. Bastos-Leite, G. R. Ridgway, C. Silveira, A. Norton, S. Reis, and K. J. Friston. Dysconnectivity within the default mode in first-episode schizophrenia: a stochastic dynamic causal modeling study with functional magnetic resonance imaging. Schizophr Bull, 41: 144–153, 2015. doi: 10.1093/schbul/sbu080.

V. Betti, S. Della Penna, F. de Pasquale, D. Mantini, L. Marzetti, G. L. Romani, and M. Corbetta. Natural scenes viewing alters the dynamics of functional connectivity in the human brain. Neuron, 79:782–797, 2013.

B. Biswal, F. Yetkin, V. Haughton, and J. Hyde. Functional connectivity in the motor cortex of resting human brain using echo-planar MRI. Magn Reson Med, 34:537–541, 1995.

K. H. Brodersen, T. M. Schofield, A. P. Leff, C. S. Ong, E. I. Lomakina, J. M. Buhmann, and K. E. Stephan. Generative embedding for model-based classification of fmri data. PLoS Comput Biol, 7:e1002079, 2011. doi: 10.1371/journal.pcbi.1002079.

V. D. Calhoun, S. M. Lawrie, J. Mourao-Miranda, and K. E. Stephan. Prediction of individual differences from neuroimaging data. Neuroimage, 145:135–136, 2017. doi: 10.1016/j.neuroimage.2016.12.012.

L. Chang, P. Gianaros, S. Manuck, A. Krishnan, and T. Wager. A sensitive and specific neural signature for picture-induced negative affect. PLoS Biol, 13:e1002180, 2015.

B. Chen, T. Xu, C. Zhou, L. Wang, N. Yang, Z. Wang, H.-M. Dong, Z. Yang, Y.-F. Zang, X.-N. Zuo, and X.-C. Weng. Individual variability and test-retest reliability revealed by ten repeated resting-state brain scans over one month. PLoS One, 10:e0144963, 2015. doi: 10.1371/journal.pone.0144963.

D. Cordes, V. M. Haughton, K. Arfanakis, G. J. Wendt, P. A. Turski, C. H. Moritz, M. A. Quigley, and M. E. Meyerand. Mapping functionally related regions of brain with functional connectivity MR imaging. Am J Neuroradiol, 21:1636–1644, 2000.

B. Da Mota, V. Fritsch, G. Varoquaux, T. Banaschewski, G. J. Barker, A. L. W. Bokde, U. Bromberg, P. Conrod, J. Gallinat, H. Garavan, J.-L. Martinot, F. Nees, T. Paus, Z. Pausova, M. Rietschel, M. N. Smolka, A. Strohle, V. Frouin, J.-B. Poline, B. Thirion, and IMAGEN consortium. Randomized parcellation based inference. Neuroimage, 89: 203–215, 2014. doi: 10.1016/j.neuroimage.2013.11.012.

G. Deco and M. L. Kringelbach. Great expectations: using whole-brain computational connectomics for understanding neuropsychiatric disorders. Neuron, 84:892–905, 2014. doi: 10.1016/j.neuron.2014.08.034.

G. Deco, V. Jirsa, and A. McIntosh. Emerging concepts for the dynamical organization of resting-state activity in the brain. Nat Rev Neurosci, 12:43–56, 2011.

A. T. Drysdale, L. Grosenick, J. Downar, K. Dunlop, F. Mansouri, Y. Meng, R. N. Fetcho, B. Zebley, D. J. Oathes, A. Etkin, A. F. Schatzberg, K. Sudheimer, J. Keller, H. S. Mayberg, F. M. Gunning, G. S. Alexopoulos, M. D. Fox, A. Pascual-Leone, H. U. Voss, B. J. Casey, M. J. Dubin, and C. Liston. Resting-state connectivity biomarkers define neurophysiological subtypes of depression. Nat Med, 23:28–38, 2017. doi: 10.1038/nm.4246.

A. Ekstrom. How and when the fMRI BOLD signal relates to underlying neural activity: the danger in dissociation. Brain Res Rev, 62:233–244, 2010.

A. K. Engel, C. Gerloff, C. C. Hilgetag, and G. Nolte. Intrinsic coupling modes: multiscale interactions in ongoing brain activity. Neuron, 80:867–886, 2013. doi: 10.1016/j.neuron. 2013.09.038.

E. Filevich, N. Lisofsky, M. Becker, O. Butler, M. Lochstet, J. Martensson, E. Wenger, U. Lindenberger, and S. Kuohn. Day2day: investigating daily variability of magnetic resonance imaging measures over half a year. BMC Neurosci, 18:65, 2017. doi: 10.1186/s12868-017-0383-y.

E. S. Finn, X. Shen, D. Scheinost, M. D. Rosenberg, J. Huang, M. M. Chun, X. Papademetris, and R. T. Constable. Functional connectome fingerprinting: identifying individuals using patterns of brain connectivity. Nat Neurosci, 18:1664–1671, 2015. doi: 10.1038/nn.4135.

E. S. Finn, D. Scheinost, D. M. Finn, X. Shen, X. Papademetris, and R. T. Constable. Can brain state be manipulated to emphasize individual differences in functional connectivity? Neuroimage, 160:140–151, 2017. doi: 10.1016/j.neuroimage.2017.03.064.

S. Frassle, K. E. Stephan, K. J. Friston, M. Steup, S. Krach, F. M. Paulus, and A. Jansen. Test-retest reliability of dynamic causal modeling for fmri. Neuroimage, 117:56–66, 2015. doi: 10.1016/j.neuroimage.2015.05.040.

P. Fries. A mechanism for cognitive dynamics: neuronal communication through neuronal coherence. Trends Cogn Sci, 9:474–480, 2005.

K. Friston. Functional and effective connectivity: A review. Brain Connect, 1:8, 2011.

K. J. Friston, L. Harrison, and W. Penny. Dynamic causal modelling. Neuroimage, 19: 1273–1302, 2003.

K. J. Friston, J. Kahan, B. Biswal, and A. Razi. A DCM for resting state fMRI. Neuroimage, 94:396–407, 2014. doi: 10.1016/j.neuroimage.2013.12.009.

M. Gilson, R. Moreno-Bote, A. Ponce-Alvarez, P. Ritter, and G. Deco. Estimation of directed effective connectivity from fMRI functional connectivity hints at asymmetries of cortical connectome. PLoS Comput Biol, 12:e1004762, 2016.

M. Gilson, G. Deco, K. Friston, P. Hagmann, D. Mantini, V. Betti, G. L. Romani, and M. Corbetta. Effective connectivity inferred from fmri transition dynamics during movie viewing points to a balanced reconfiguration of cortical interactions. NeuroImage, 2017. doi: https://doi.org/10.10167j.neuroimage.2017.09.061.

R. Goebel, A. Roebroeck, D. Kim, and E. Formisano. Investigating directed cortical interactions in time-resolved fMRI data using vector autoregressive modeling and Granger causality mapping. Magn Reson Imaging, 21:1251–1261, 2003.

J. Gonzalez-Castillo and P. A. Bandettini. Task-based dynamic functional connectivity: Recent findings and open questions. Neuroimage, 2017. doi: 10.1016/j.neuroimage.2017.08.006.

J. Gonzalez-Castillo, C. W. Hoy, D. A. Handwerker, M. E. Robinson, L. C. Buchanan, Z. S. Saad, and P. A. Bandettini. Tracking ongoing cognition in individuals using brief, whole-brain functional connectivity patterns. Proc Natl Acad Sci USA, 112:8762–8767, 2015. doi: 10.1073/pnas.1501242112.

E. M. Gordon, T. O. Laumann, A. W. Gilmore, D. J. Newbold, D. J. Greene, J. J. Berg, M. Ortega, C. Hoyt-Drazen, C. Gratton, H. Sun, J. M. Hampton, R. S. Coalson, A. L. Nguyen, K. B. McDermott, J. S. Shimony, A. Z. Snyder, B. L. Schlaggar, S. E. Petersen, S. M. Nelson, and N. U. F. Dosenbach. Precision functional mapping of individual human brains. Neuron, 95:791–807, 2017. doi: 10.1016/j.neuron.2017.07.011.

M. Greicius. Resting-state functional connectivity in neuropsychiatric disorders. Curr Opin Neurol, 21:424–430, 2008. doi: 10.1097/WCO.0b013e328306f2c5.

I. Guyon, J. Weston, S. Barnhill, and V. Vapnik. Gene selection for cancer classification using support vector machines. Machine Learning, 46(1):389–422, 2002. doi: 10.1023/A:1012487302797.

P. Hagmann, L. Cammoun, X. Gigandet, R. Meuli, C. J. Honey, V. J. Wedeen, and O. Sporns. Mapping the structural core of human cerebral cortex. PLoS Biol, 6:e159, 2008.

B. J. He. Scale-free properties of the functional magnetic resonance imaging signal during rest and task. J Neurosci, 31:13786–13795, 2011.

J. F. Hipp, A. K. Engel, and M. Siegel. Oscillatory synchronization in large-scale cortical networks predicts perception. Neuron, 69:387–396, 2011.

M. J. Hoptman, X.-N. Zuo, D. D’Angelo, C. J. Mauro, P. D. Butler, M. P. Milham, and D. C. Javitt. Decreased interhemispheric coordination in schizophrenia: a resting state fmri study. Schizophr Res, 141:1–7, 2012. doi: 10.1016/j.schres.2012.07.027.

G. Hughes. On the mean accuracy of statistical pattern recognizers. IEEE Transactions on Information Theory, 14:55–63, 1968. doi: 10.1109/TIT.1968.1054102.

T. Kaufmann, D. Alnæs, N. T. Doan, C. L. Brandt, O. A. Andreassen, and L. T. Westlye. Delayed stabilization and individualization in connectome development are related to psychiatric disorders. Nat Neurosci, 20:513–515, 2017. doi: 10.1038/nn.4511.

S. Kurth, E. Moyse, M. A. Bahri, E. Salmon, and C. Bastin. Recognition of personally familiar faces and functional connectivity in alzheimer’s disease. Cortex, 67:59–73, 2015. doi: 10.1016/j.cortex.2015.03.013.

B. Li, X. Wang, S. Yao, D. Hu, and K. Friston. Task-dependent modulation of effective connectivity within the default mode network. Front Psychol, 3:206, 2012. doi: 10.3389/fpsyg.2012.00206.

D. Mantini, U. Hasson, V. Betti, M. G. Perrucci, G. L. Romani, M. Corbetta, G. A. Orban, and W. Vanduffel. Interspecies activity correlations reveal functional correspondence between monkey and human brain areas. Nat Methods, 9:277–282, 2012.

P. M. Matthews and A. Hampshire. Clinical concepts emerging from fmri functional connectomics. Neuron, 91:511–528, 2016. doi: 10.1016/j.neuron.2016.07.031.

A. Messe, D. Rudrauf, H. Benali, and G. Marrelec. Relating structure and function in the human brain: relative contributions of anatomy, stationary dynamics, and non-stationarities. PLoS COmput Biol, 10:e1003530, 2014.

O. Miranda-Dominguez, B. D. Mills, S. D. Carpenter, K. A. Grant, C. D. Kroenke, J. T. Nigg, and D. A. Fair. Connectotyping: model based fingerprinting of the functional connectome. PLoS One, 9:e111048, 2014. doi: 10.1371/journal.pone.0111048.

A. Mitra, A. Z. Snyder, E. Tagliazucchi, H. Laufs, and M. Raichle. Propagated infra-slow intrinsic brain activity reorganizes across wake and slow wave sleep. Elife, 4, 2015.

S. Mueller, D. Wang, M. D. Fox, R. Pan, J. Lu, K. Li, W. Sun, R. L. Buckner, and H. Liu. Reliability correction for functional connectivity: Theory and implementation. Hum Brain Mapp, 36:4664–4680, 2015. doi: 10.1002/hbm.22947.

M. Pannunzi, R. Hindriks, R. G. Bettinardi, E. Wenger, N. Lisofsky, J. Martensson, O. Butler, E. Filevich, M. Becker, M. Lochstet, S. Kuhn, and G. Deco. Resting-state fmri correlations: From link-wise unreliability to whole brain stability. Neuroimage, 157:250–262, 2017. doi: 10.1016/j.neuroimage.2017.06.006.

M. G. Preti, T. A. Bolton, and D. Van De Ville. The dynamic functional connectome: State-of-the-art and perspectives. Neuroimage, 160:41–54, 2017. doi: 10.1016/j.neuroimage.2016.12.061.

M. Rahim, B. Thirion, D. Bzdok, I. Buvat, and G. Varoquaux. Joint prediction of multiple scores captures better individual traits from brain images. Neuroimage, 158:145–154, 2017. doi: 10.1016/j.neuroimage.2017.06.072.

M. E. Raichle, A. M. MacLeod, A. Z. Snyder, W. J. Powers, D. A. Gusnard, and G. L. Shulman. A default mode of brain function. Proc Natl Acad Sci USA, 98:676–682, 2001. doi: 10.1073/pnas.98.2.676.

J. Rissman and A. D. Wagner. Distributed representations in memory: insights from functional brain imaging. Annu Rev Psychol, 63:101–128, 2012. doi: 10.1146/annurev-psych-120710-100344.

P. Rousseeuw. Silhouettes: A graphical aid to the interpretation and validation of cluster analysis. Journal of Computational and Applied Mathematics, 20:53–65, 1987. doi: https://doi.org/10.1016/0377-0427(87)90125-7.

M. Senden, N. Reuter, M. P. van den Heuvel, R. Goebel, G. Deco, and M. Gilson. Task-related effective connectivity reveals that the cortical rich club gates cortex-wide communication. Hum Brain Mapp, 2017. doi: 10.1002/hbm.23913.

D. V. Sheehan, Y. Lecrubier, K. H. Sheehan, P. Amorim, J. Janavs, E. Weiller, T. Hergueta, R. Baker, and G. C. Dunbar. The mini-international neuropsychiatric interview (m.i.n.i.): the development and validation of a structured diagnostic psychiatric interview for dsm-iv and icd-10. J Clin Psychiatry, 59:22–33, 1998.

Z. Shehzad, A. M. C. Kelly, P. T. Reiss, D. G. Gee, K. Gotimer, L. Q. Uddin, S. H. Lee, D. S. Margulies, A. K. Roy, B. B. Biswal, E. Petkova, F. X. Castellanos, and M. P. Milham. The resting brain: unconstrained yet reliable. Cereb Cortex, 19:2209–2229, 2009. doi: 10.1093/cercor/bhn256.

H. Shen. Neuroscience: Tuning the brain. Nature, 507:290–292, 2014. doi: 10.1038/507290a.

K. Stephan, L. Harrison, W. Penny, and K. Friston. Biophysical models of fMRI responses. Curr Opin Neurol, 14:629–635, 2004.

K. E. Stephan and C. Mathys. Computational approaches to psychiatry. Curr Opin Neurobiol, 25:85–92, 2014. doi: 10.1016/j.conb.2013.12.007.

N. Tzourio-Mazoyer, B. Landeau, D. Papathanassiou, F. Crivello, O. Etard, N. Delcroix, B. Mazoyer, and M. Joliot. Automated anatomical labeling of activations in spm using a macroscopic anatomical parcellation of the MNI MRI single-subject brain. Neuroimage, 15:273–289, 2002.

T. Vanderwal, J. Eilbott, E. S. Finn, R. C. Craddock, A. Turnbull, and F. X. Castellanos. Individual differences in functional connectivity during naturalistic viewing conditions. Neuroimage, 157:521–530, 2017. doi: 10.1016/j.neuroimage.2017.06.027.

G. Varoquaux, P. R. Raamana, D. A. Engemann, A. Hoyos-Idrobo, Y. Schwartz, and B. Thirion. Assessing and tuning brain decoders: Cross-validation, caveats, and guidelines. Neuroimage, 145:166–179, 2017. doi: 10.1016/j.neuroimage.2016.10.038.

C.-W. Woo, L. J. Chang, M. A. Lindquist, and T. D. Wager. Building better biomarkers: brain models in translational neuroimaging. Nat Neurosci, 20:365–377, 2017. doi: 10.1038/ nn.4478.

H. Xie, V. D. Calhoun, J. Gonzalez-Castillo, E. Damaraju, R. Miller, P. A. Bandettini, and S. Mitra. Whole-brain connectivity dynamics reflect both task-specific and individual-specific modulation: A multitask study. Neuroimage, 2017. doi: 10.1016/j.neuroimage.2017.05.050.

N. Yahata, K. Kasai, and M. Kawato. Computational neuroscience approach to biomarkers and treatments for mental disorders. Psychiatry Clin Neurosci, 71:215–237, 2017. doi: 10.1111/pcn.12502.

C.-G. Yan, X.-D. Wang, X.-N. Zuo, and Y.-F. Zang. Dpabi: Data processing & analysis for (resting-state) brain imaging. Neuroinformatics, 14:339–351, 2016. doi: 10.1007/s12021-016-9299-4.

X.-N. Zuo, J. S. Anderson, P. Bellec, R. M. Birn, B. B. Biswal, J. Blautzik, J. C. S. Breitner, R. L. Buckner, V. D. Calhoun, F. X. Castellanos, A. Chen, B. Chen, J. Chen, X. Chen, S. J. Colcombe, W. Courtney, R. C. Craddock, A. Di Martino, H.-M. Dong, X. Fu, Q. Gong, K. J. Gorgolewski, Y. Han, Y. He, Y. He, E. Ho, A. Holmes, X.-H. Hou, J. Huckins, T. Jiang, Y. Jiang, W. Kelley, C. Kelly, M. King, S. M. LaConte, J. E. Lainhart, X. Lei, H.-J. Li, K. Li, K. Li, Q. Lin, D. Liu, J. Liu, X. Liu, Y. Liu, G. Lu, J. Lu, B. Luna, J. Luo, D. Lurie, Y. Mao, D. S. Margulies, A. R. Mayer, T. Meindl, M. E. Meyerand, W. Nan, J. A. Nielsen, D. O’Connor, D. Paulsen, V. Prabhakaran, Z. Qi, J. Qiu, C. Shao, Z. Shehzad, W. Tang, A. Villringer, H. Wang, K. Wang, D. Wei, G.-X. Wei, X.-C. Weng, X. Wu, T. Xu, N. Yang, Z. Yang, Y.-F. Zang, L. Zhang, Q. Zhang, Z. Zhang, Z. Zhang, K. Zhao, Z. Zhen, Y. Zhou, X.-T. Zhu, and M. P. Milham. An open science resource for establishing reliability and reproducibility in functional connectomics. Sci Data, 1:140049, 2014. doi: 10.1038/sdata.2014.49.

